# NEURON-SPECIFIC CHROMOSOMAL MEGADOMAIN ORGANIZATION IS ADAPTIVE TO RECENT RETROTRANSPOSON EXPANSIONS

**DOI:** 10.1101/2021.08.30.458289

**Authors:** Sandhya Chandrasekaran, Sergio Espeso-Gil, Yong-Hwee Eddie Loh, Behnam Javidfar, Bibi Kassim, Yuhao Dong, Yueyan Zhu, Lucy K. Bicks, Prashanth Rajarajan, Cyril J. Peter, Esperanza Agullo-Pascual, Marina Iskhakova, Molly Estill, Li Shen, Yan Jiang, Schahram Akbarian

## Abstract

Here, we mapped cell-type specific chromatin domain organization in adult mouse cerebral cortex and report strong enrichment of *Endogenous Retrovirus 2 (ERV2)* repeat sequences in the neuron-specific heterochromatic ‘B_2_^NeuN+^’ megabase-scaling subcompartment. Comparative chromosomal conformation mapping in *Mus spretus* and *Mus musculus* revealed neuron-specific reconfigurations tracking recent ERV2 retrotransposon expansions in the murine germline, with significantly higher B_2_ megadomain contact frequencies at sites with ongoing ERV2 insertions in *Mus musculus*. Ablation of the retrotransposon silencer *Kmt1e/Setdb1* triggered B2 megadomain disintegration and rewiring with open chromatin domains enriched for cellular stress response genes, along with severe neuroinflammation and proviral assembly of ERV2/Intracisternal-A-Particles (IAPs) infiltrating dendrites and spines. We conclude that neuronal megadomain architectures include evolutionarily adaptive heterochromatic organization which, upon perturbation, unleashes ERV proviruses with strong tropism within mature neurons.

---

Repeat-rich sequence blocks, considered major determinants for 3D folding and structural genome organization in the cell nucleus in all higher eukaryotes, are critically involved in a wide range of genomic functions, from lineage-specific gene expression programs in fungi (*1*) to X-inactivation in early mammalian development (*2*). Repetitive DNA may also be important for spatial genome organization in the brain. For example, monogenic neurodegenerative and neurodevelopmental diseases could result from abnormal locus-specific expansion of short-tandem repeats (STR) at the periphery of topologically-associating domains (TADs), a type of conformation defined by chromosomal loop extrusions normally constrained by strong boundary elements at TAD peripheries (*3*). However, the relationship between the 3D genome (3DG) and DNA repeat organization in brain cells, including potential implications for neuronal health and function, remains unexplored. Here, we show that megabase-scale chromatin domain organization in adult mouse cerebral cortex is linked, in highly cell type-specific fashion, to multiple retrotransposon superfamilies, comprising the vast majority of ‘mobile’ DNA elements in the murine genome (*4*). We identify a neuronal megadomain subtype for which species-specific interaction frequencies track the dramatic reconfiguration of the retrotransposon landscape in *Mus musculus*-derived inbred lines, primarily due to ongoing germline expansions of ERV *Endogenous Retrovirus* elements. Neuronal deficiency for *Kmt1e/Setdb1* histone methyltransferase, critical to the KMT1E-KAP1-Zinc finger and retrotransposon silencer complex (*5, 6*), triggered massive megabase-scale disintegration and rewiring of chromosomal interactions among chromatin domains anchored in ERV-rich genomic loci. This was associated with retrotransposon un-silencing and severe neuroinflammation and activation of cellular stress genes, intriguingly in close physical proximity to ERV-enriched megadomains, with the endomembrane system of susceptible Setdb1-deficient neurons hijacked for provirus assembly generating provirus-like particles. Our findings provide the first example of how 3DG compartmentalization in the mature mouse brain is critically shaped by mobile DNA elements in strictly cell-type fashion, uncovering a distinct heterochromatic regulome in neurons which, upon perturbation, could robustly unleash ERV proviruses.

## Results

### Megadomain organization in adult mouse cerebral cortex

To study neuronal megadomain organization in adult mouse cerebral cortex, we prepared *in situ* Hi-C libraries from neuronal (NeuN+), and for comparison, non-neuronal (NeuN−) nuclei from N=4 (2F/2M) mice) and sorted HiC Pro (v2.9) chimeric reads from autosomal sequences by k-means clustering at 250kb resolution (Figure 1A, Table S1) (*7*). We identified 4 large chromosomal subcompartments in each population, comprising 2.46Gb of sequence, or >99.9% of the autosomal murine reference genome *mm10* -- in NeuN+ (*range*: 104- 1460Mb/subcompartment) and in NeuN− (range: 95-1452Mb/subcompartment) (Figures S1, S2, Tables S2, S3), and assigned identifiers to each first according to minority (A) or majority (B) fractional concordance with heterochromatic, nuclear lamina-associated domain (LAD) sequences, and then numbered based on decreasing size (Figure 1B). Similar loci comprised subcompartments within NeuN+ and NeuN− nuclei, with the notable exception of loci within B2; B_2_^NeuN−^ loci included substantial loci from both B_1_^NeuN+^ and B_2_^NeuN+^. This would suggest that in NeuN+, B_2_^NeuN+^ interact in a unique manner as compared to NeuN−. Indeed, average *trans* interaction frequencies among B_2_^NeuN+^ and B_2_^NeuN−^ markedly differed, in contrast to otherwise similar interaction profiles for the remaining subcompartments (Figure 1C). Chromatin profiling (mouseENCODE (*8*)) identified two subcompartments in NeuN+ (*A_1_^NeuN+^* and *A_2_^NeuN+^)* and NeuN− (*A_1_^NeuN−^* and *A_2_^NeuN−^)* with high fractional concordances for enhancers and chromatin marks broadly associated with actual or potential transcription (Figure 1D). Uniquely, B_2_^NeuN+^ revealed a high (60.1%) fractional concordance with the H3K9me3 repressive histone mark, unseen in B_2_^NeuN−^ (Figure 1D). In fact, neuronal B_2_^NeuN+^ megadomain sequences showed a highly specific, significant association (p=3.36×10^−41^, Fisher’s 2×2 exact test) with adult cortex NeuN+ H3K9me3, but not with active histone marks (Figures 1E, 1F, Table S4) (*9*), also unseen in NeuN− nuclei. In addition, we noticed that 89.6Mb (86.2%) of B_2_^NeuN+^ sequences anchored the vast majority of neuron-specific inter-autosomal contacts (Homer v4.8, 1Mb resolution, p<10^−50^) were not shared with non-neuronal cells (Figure S3, Tables S5, S6). As a result, *trans* contact maps sharply diverged by cell type, with striking consistency across the 4 animals (Pearson’s r range: NeuN+ [0.81 – 0.95] & NeuN− [0.95 – 0.99]) (Figure 1G). Interestingly, neuron-specific *trans* interactions predominantly involved B_2_^NeuN+^ loci, while non-neuron-specific *trans* interactions primarily involved A_1_^NeuN−^ and A_2_^NeuN−^ (Figure 1H, Figure S4). Notably, several gene clusters, including protocadherins, vomeronasal receptors, and cytochrome p450 enzymes, are encompassed within B_2_^NeuN+^ megadomain interactions (Figure S5). Robust interchromosomal Hi-C contacts of olfactory receptor gene clusters were previously reported in olfactory sensory neurons (*10*).

**Figure 1:**
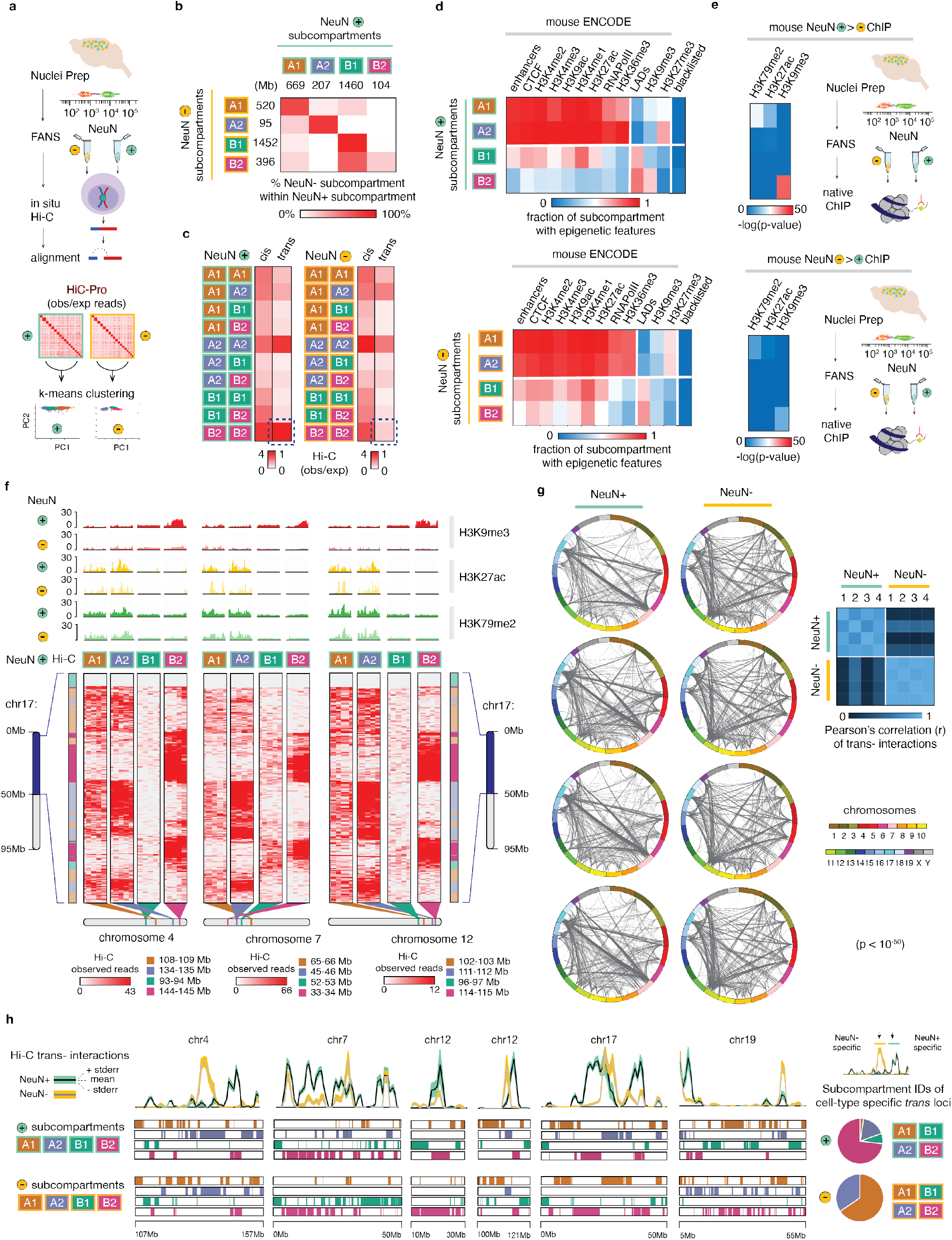
(A) (Left) Workflow. Fluorescence activated nuclei sorting (FANS) was performed on dissected adult mouse cortical tissue (n=4, 2F/2M), followed by *in situ* Hi-C on intact NeuN+ and NeuN− nuclei. Genome alignment (HiC- Pro (v2.9)) facilitated by split-mapping at the known chimeric ligation junctions to generate genome-wide pairwise contact maps. Hi-C read counts for NeuN+ and NeuN− (observed/expected) were then independently piped into a k-means clustering algorithm, ultimately generating four clusters representative of chromatin subcompartments in each population. (B) Correspondence of the NeuN+ and NeuN− determined subcompartments; genome extents of each subcompartment as indicated. Percent overlap of coordinates from each NeuN− subcompartment with each of the NeuN+ subcompartments represented on indicated color scale. (C) Heatmap of mean Hi-C (observed/expected) read counts for NeuN+ (left) and NeuN− (right) between loci comprising the designated subcompartments; 250kb resolution. (D) Heatmap summarizing the characterization of determined clusters as ‘A’ (active) or ‘B’ (inactive) with lamin-associated domains (LADs), enhancer coordinates, and mENCODE ChIP- Seq data for RNAPolII, CTCF, and several histone modifications as indicated, and mENCODE blacklisted sequences in NeuN+ (top) and NeuN− (bottom). The fraction of 250kb bins comprising the cluster overlapping these genome tracks is indicated on a scale of 0 (blue) to 1 (red). (E) Enrichment heatmap of differential NeuN+ and NeuN− H3K79me2 (n=2), H3K27ac (n=3), and H3K9me3 (n=4) tagged sequences with loci comprising each of the four subcompartments. Top: NeuN+ histone modification enrichment, diffReps (p<0.001). Bottom: NeuN− histone modification enrichment, diffReps (p<0.001). (F) Representative NeuN+ long-range contact patterns (250kb resolution) for multiple genomic intervals comprising each chromatin subcompartment. Normalized NeuN+ and NeuN− histone modification profiles tracks included. (G) Individual Circos plots from four biological replicates for NeuN+ (left) and NeuN− (right) trans-chromosomal contacts (1Mb resolution, HOMER (v4.8), p<10^− 50^). NeuN+ and NeuN− Circos plots aligned vertically originate from the same brain tissue. (Right) Pearson’s correlation heatmap of loci involved in trans-interactions in NeuN+ and NeuN− across replicates. (H) (Left) Hi-C pairwise *trans* contact maps for NeuN+ (green) and NeuN− (orange) across several chromosomes. NeuN+ (n=4) mean *trans* interactions in black, standard error in green; NeuN− (n=4) mean *trans* interactions in gray, standard error in orange. Subcompartment designations in NeuN+ (top) and NeuN− (bottom) included. (Right) Top: Percent of NeuN+ specific *trans* interactions (n=88 1Mb loci, defined as 250kb loci with a mean of >1 significant *trans* interaction in NeuN+ (n=4) and <1 in NeuN− (n=4), as categorized by NeuN+ subcompartment. Bottom: Percent of NeuN− specific *trans* interactions (n=197 1Mb loci, defined as 250kb loci with a mean of >1 significant *trans* interaction in NeuN− (n=4) and <1 in NeuN+ (n=4), as categorized by NeuN− subcompartment.

Transposable elements, simple repeats and other types of repetitive DNA comprise up to 45% of the mouse genome (*11*). Therefore, we asked whether our megadomain subcompartments show repeat-specific enrichment. We screened our four subcompartments against 10 broad classes of repeat categories (Table S7). Indeed, B_2_^NeuN+^ showed a significant (p= 2.13×10^−35^, Fisher’s 2×2 exact test) enrichment for hotspots (99^th^ percentile densities) of the *Endogenous Retrovirus* subtype *ERV2* (Figure 2A). The *ERVs* are a vertebrate-specific RNA-based (’copy-and-paste’) retrotransposon family defined by long-terminal repeats (LTRs) flanking the core *gag-pol-env* viral coding domains (*12, 13*); as a class, ERVs colonize approximately 10% of the mouse genome (ERV2 specifically comprises 3.14%) (*14*). Remarkably, none of the remaining neuronal subcompartments and none of the non-neuronal subcompartments showed *ERV2* enrichment. However, both A_2_^NeuN+^ and A_2_^NeuN−^ showed significant enrichment (p= 2.47×10^−55^, Fisher’s 2×2 exact test) for *SINE* (*Short-Interspersed Nucleotide Elements)* non-autonomous retroelements (*12*) previously linked to domain boundaries and inter-chromosomal interactions in some metazoan systems (*15–17*), serving as CTCF binding scaffolds (Figure S6). Furthermore, the A_1_ subcompartment in both cell types was significantly enriched for the largely fossilized ERV3 TEs. Furthermore, *LINE* (Long-interspersed Nucleotide Element) TE showed strong enrichment for the B1 compartment for both cell types. Thus, each major type of TE shows unique, subcompartment-specific enrichment patterns in brain, with neuron-specific B_2_^NeuN+^ megadomains tagged with high levels of H3K9me3 and enriched for ERV2 elements.

**Figure 2:**
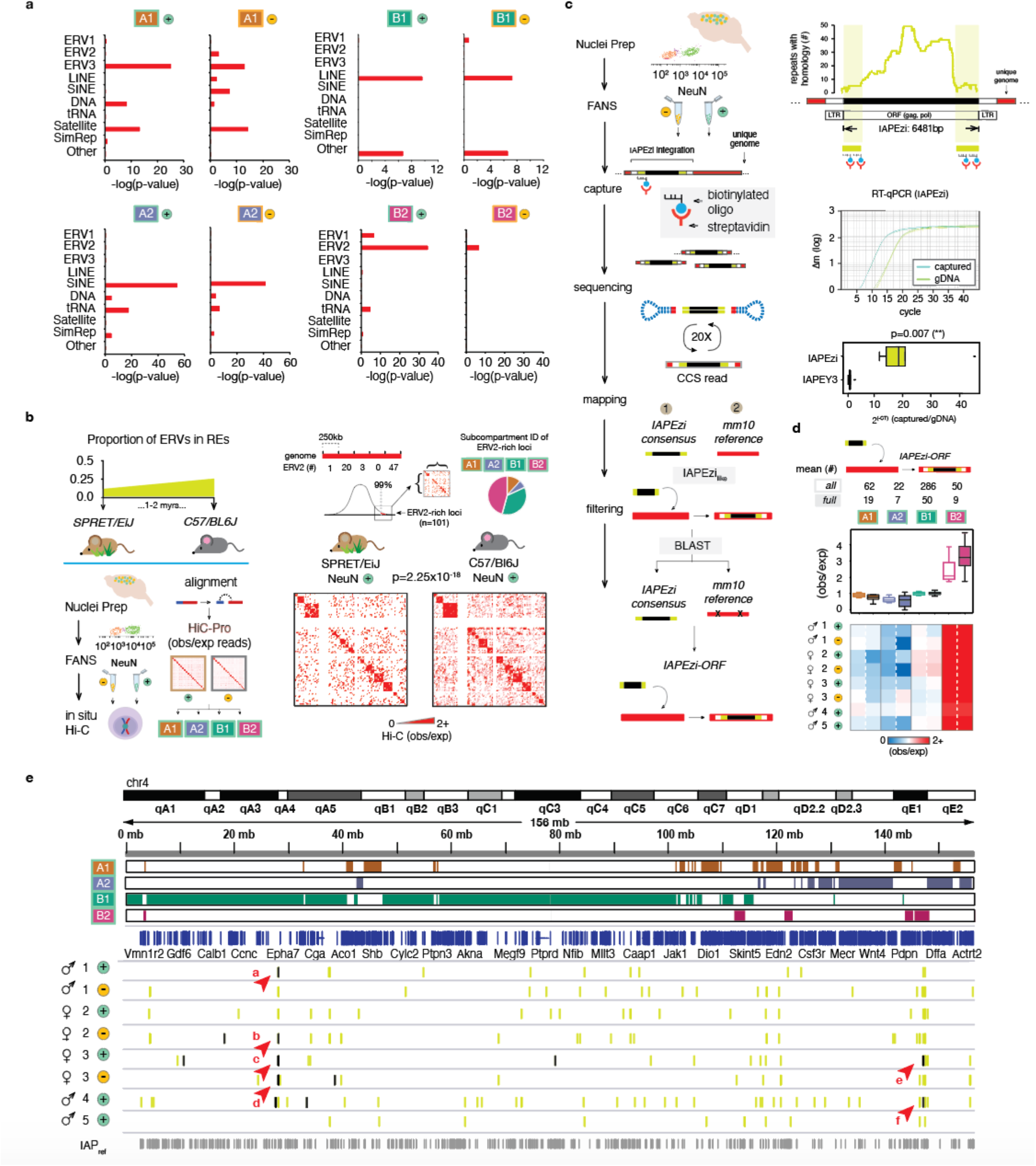
(A) Repetitive element (RE) associations with loci comprising NeuN+ (left) and NeuN− (right) subcompartments. REs (Repbase) classified as indicated. -Log(p-values) displayed as red bars (Fisher’s 2×2 test, one-tailed). (B) (Left) Schematic of workflow. (Top) Proportion of ERVs within REs in SPRET/EiJ and C57/BL6J; adapted from(*18*). (Bottom) FANS on dissected adult SPRET/EiJ and C57/BL6J mouse cortical tissue (n=4, 2M/2F), followed by *in situ* Hi-C on the sorted intact NeuN+ and NeuN− nuclei, aligned to the genome using HiC-Pro (v2.9) to generate Hi-C pairwise interaction matrices. (Right) Summary matrix of NeuN+ Hi-C values (observed//expected) denoting interaction frequencies between genomic loci harboring high densities of ERV2 (>99^th^ percentile, 250kb resolution) in C57/BL6J mice as compared to SPRET/EiJ (p=2.25×10^−18^), Wilcoxon sum- rank test, two-sided, paired). Pie chart denotes the subcompartment distribution of the analyzed ERV2-rich loci. (C) (Left) Schematic of PacBio SMRT circular consensus sequence (CCS) long-read sequencing workflow. *De novo* reads identified by evaluating BLAST bit scores between alignment to the IAPEzi consensus and the IAP- masked reference genome. (Right) (Top) PacBio biotinylated oligonucleotide probe design. The probe sequences are the most divergent regions of IAPEzi across other repetitive elements with detectable homology (<10%, *dfam.org*). (Bottom) Real time qPCR plot (n=6) of genomic DNA vs. IAPEzi-captured DNA. Representative qPCR trace included. (D) Subcompartment observed/expected heatmap for 8 (Mus musculus, 3 C57/BL6, 5 129S1) mouse cortical NeuN+ and NeuN− samples IAPEzi *de novo* integration sites, as indicated. Full *de novo* reads refer to *de novo* reads with >90% (5833/6481bp) match with the IAPEzi consensus sequence length (*dfam.org*). (E) Genome browser shot of chr4 with *de novo* IAPEzi insertions for each of 8 profiles samples. Subcompartments designations as indicated. Arrows point to representative full-length *de novo* insertions in close genomic range of each other across different samples. Specific coordinates corresponding to all (green) vs. full (black) *de novo* in Supplementary Table 8).

Of note, a subset of ERVs with preserved retrotransposon capacity, including intracisternal A particles (IAPs), continue to invade the murine germline, accelerating their expansion within *Mus musculus*-derived inbred lines (*18*). As a result, IAPs comprise >80% of all ERVs genome-wide in C57/BL6J laboratory inbred mice, in stark contrast to the very low (<10%) phylogenetic conservation of those IAPs in the genomes of wild-derived *SPRET/EiJ* mice (*18*). We wondered whether inter-chromosomal contacts, comprising the defining feature of the B_2_^NeuN*+*^ subcompartment, occur in reduced frequencies in *SPRET/EiJ* vs. *C57/BL6J* neurons. To explore, we prepared Hi-C libraries from sorted adult cortex NeuN+ and NeuN− nuclei of both mouse strains in parallel (n=2 (1F/1M/strain)) (Figure 2B). Hotspots of ERV2 repeats (n=101 250kb bins), defined as >99^th^ percentile densities of ERV2 in the *mm10* reference genome, interacted less frequently in SPRET/EiJ versus C57/BL6J NeuN+ (p=2.25×10^−18^); in contrast, interaction frequencies at these loci did not significantly differ in NeuN− nuclei across the two strains (Figure S7), supporting a role for ERV2 densities in shaping the 3D NeuN+ genome. We conclude that B_2_^NeuN+^ megadomain conformations in mature cortical neurons show ‘dosage-sensitivity’ for phylogenetically young ERV2 sequences.

We suspected that many IAP integration sites in our profiled mouse brains are, in fact, not represented in the reference genome due to a variety of factors including residual genetic contribution from non-C57/BL6J lines, genetic drift, or somatic retrotransposition in stem cells and progenitor cells during early embryogenesis or in the developing brain. To explore whether the observed IAP/ERV2 enrichment of the B_2_^NeuN+^ subcompartment is preserved beyond the retroelement sequences annotated in *mm10*, we used biotinylated oligonucleotides to capture 10-15kb sized DNA fragments carrying phylogenetically young and retrotransposition-competent IAP subtype *IAPEzi* (6481bp) (*14, 19, 20*) together with surrounding flanking sequences for genomic annotation (Figure 2C). We ran single molecule PacBio SMRT-sequencing (SMRT-seq) on captured DNA of sorted NeuN+ neuronal and, separately, NeuN− non-neuronal nuclei collected from adult cerebral cortex of three *C57Bl6/J* mice and two additional animals of mixed (predominantly C57Bl6/J) genetic background, generating 2.64- 4.70×10^6^ high fidelity (HiFi, >99.9% accuracy) circular consensus sequences (CCS) for each of the 8 cell-type specific samples (Table S8). For each sample, the overwhelming majority of *IAPEzi* sequences (83-94%) expectedly annotated to IAPEzi *GRCm38/mm10* sites (Repeatmasker). However, each of the 8 profiled samples identified 209-684 proviral (non-solo-LTR) *IAPEzi* integration sites not found in *mm10*, including 2-44 sites/animal harboring full-length IAPEzi. Strikingly, proviral (non-solo LTR) sequences, including full-length *non-mm10* IAPEzi, showed a significant 2.5- to 4-fold enrichment for B_2_^NeuN+^ sequences compared to the remaining subcompartments (B_2_^all^ vs. B_2_^full^, paired Wilcoxon sum-rank test, p=0.0391 [*]) (Figure 2D, 2E, Figure S8, Table S9). We conclude that full-length, potentially retrotransposition-competent IAP elements continue to preferentially insert into genomic loci ‘destined’ to assemble as B_2_^NeuN+^ subcompartment.

### Neuronal megadomain reorganization upon ablation of Setdb1 methyltransferase

Next, we generated *CamK-Cre^+^,Setdb1^2lox/2lox^* conditional mutant mice for neuron-specific ablation of the *Kmt1e/Setdb1* histone methyltransferase, an essential regulator for ERV repression in stem cells and somatic tissues (*5*) and part of a repressor complex with SMARCAD1 chromatin remodeler and KRAB-associated protein 1 (KAP1)-KRAB zinc finger proteins (*6*). To test whether *Setdb1* could exert a broad regulatory footprint of *Setdb1* on ERV-rich megadomains, including compartmental organization and connectivity of neuronal B_2_^NeuN+^, we generated two NeuN+ libraries to supplement the two previously published cell-type Hi-C chromosomal contacts maps from adult *CamK-Cre^+^,Setdb1^2lox/2lox^* with littermate controls (CamK-Cre^−^, Setdb1*^2lox/^*^2lox^) (N=4/group (2M/2F) (Figure 3A, Figure S9). Differential analysis by genotype (DESeq2, 1Mb bins, FDR p_adj_ < 0.05) revealed widespread megadomain rewiring in mutant neurons -- N=446 significant decreases in pairwise *trans* interactions, the vast majority (60.76%) of which reflected loss of B_2_-B_2_ *trans*, and another N=196 pairwise increases, of which 72.45% represented gains in B_1_^NeuN+^-B_2_^NeuN+^ *trans* (Figure 3B, Tables S10, S11). Furthermore, mutant neurons showed significant shifts in intra-chromosomal *cis* Hi-C interactomes, overwhelmingly driven by ERV-enriched subcompartment sequences (42.4% of N=4066 representing involving B_2_^NeuN+^ contacts, DESeq2, 1Mb bins, FDR p_adj_ < 0.05). This included significant losses in cross-compartmental A_2_^NeuN+^-B_2_^NeuN+^ contacts (N=3.79 cis contacts/per 10Mb A_2_^NeuN+^), representing a 2-fold enrichment as compared to A_1_^NeuN+^-B_2_^NeuN+^ contacts (N=2.05 cis contacts/per 10Mb A_1_^NeuN+^) (Figure 3B, Table S11). Of note, this biased disintegration of A_2_^NeuN+^-B_2_^NeuN+^ *cis* contacts after neuronal *Setdb1* ablation was highly specific because only 0.4 A_2_^NeuN+^_-_B_2_^NeuN+^ cis contacts/10Mb A_2_^NeuN+^ showed mutant-specific gain, while the number of A_1_^NeuN+^-B_2_^NeuN+^ contacts gained was 3.01 cis contacts/10Mb A_1_^NeuN+^.

**Figure 3:**
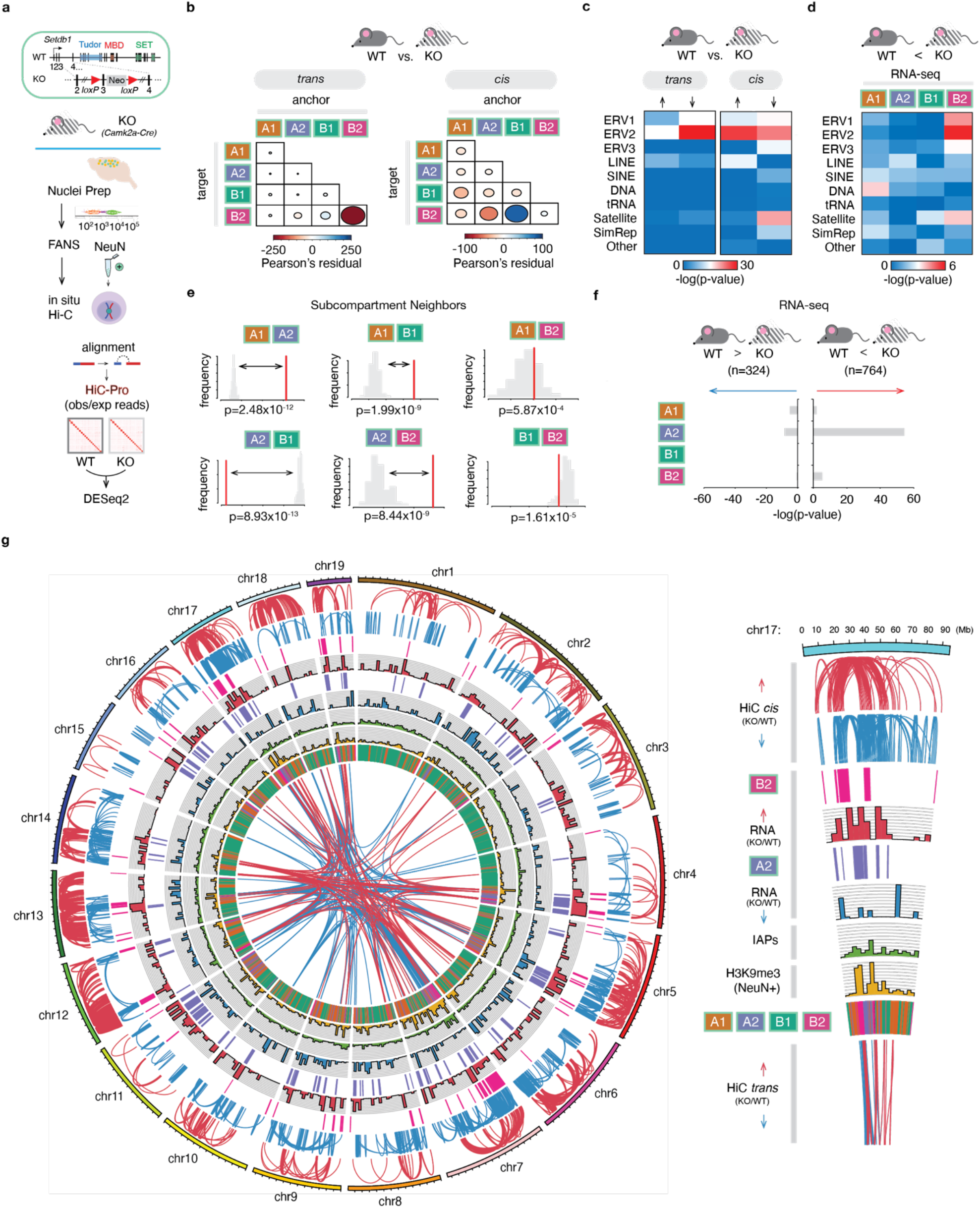
(A) Workflow. (Top) Representation of conditional Setdb1 ablation (KO), adapted from (21). (Bottom) FANS was performed on dissected adult KO mouse cortical tissue (n=4, 2F/2M), followed by in situ Hi-C on intact NeuN+ nuclei. Genome alignment (HiC-Pro (v2.9)) facilitated by split-mapping at the known chimeric ligation junctions to generate genome-wide pairwise contact maps. (B) Differential Hi-C trans (left) and cis (right) interactions in KO vs. WT (n=4 (2F/2M)/genotype (padj<0.05) (1Mb resolution). The difference in Pearson’s residuals (observed/expected) (r) between the distribution of increased interactions and decreased interactions, with size of each ellipse representing rabsolute (|r|), and color of each ellipse representing r. (C) Genome-wide associations of repetitive element families with loci involved in the altered trans and cis interactions in KO vs. WT. (D) Genome-wide associations of differentially increased RNA-Seq (n=6 (3F/3M)/genotype) of repetitive element families with subcompartment loci. (C, D) -Log(p-values) represent outcomes from Fisher’s 2×2 testing (one-tailed). (E) Proximity analyses of subcompartment loci in cis. Relative frequencies of contiguous subcompartment block neighbors in the reference genome displayed in red, while the distribution of 100 permutations (regionER) in a random genome with equivalent proportions of such blocks displayed in gray. (F) Genome-wide associations of differential (both increased and decreased) RNA-Seq (n=6 (3F/3M)/genotype) transcripts by subcompartment (- log(p-values), Fisher’s 2×2 testing (one-tailed)). (G) (Left) Circos plot representation of multiple epigenomic features in KO vs. WT NeuN+. (Right) Pie slice displays chr17 with associated legend for individual tracks applicable to the entire Circos plot, including (to top) HiC trans interaction changes (KO vs. WT), H3K9me3 enrichment, IAP densities, RNA changes (KO vs. WT), and HiC cis interactions changes (KO vs. WT).

Importantly, both *trans* and *cis* Hi-C alterations upon *Setdb1* ablation were significantly associated with ERV2 hotspots (Figure 3C). To examine whether these changes in compartment-specific chromosomal conformations are associated with transposon un-silencing, we RNA-seq profiled the cortical transcriptome of adult mutant and control mice (N=6/group) (Table S12). Importantly, comparison of subcompartment-specific expression of ERV2s and other repeat classes in mutant and control cortex revealed that increased ERV2 expression in *Setdb1*-deficient mice primarily originated within B_2_^NeuN+^, while the remaining subcompartments showed very minimal alterations in transposon expression in comparison to control animals (Figure 3D). Furthermore, expression of the 764 unique autosomal gene transcripts significantly up-regulated in mutant cortex (DESeq2, Tophat, FDR 0.05) largely originated in A_2_^NeuN+^ (Fisher’s 2×2 test, A_2_ p=3.56×10^−55^), the A subcompartment that preferentially lost *cis* contacts with B_2_^NeuN+^ (as mentioned above) and also most frequently neighbors B_2_^NeuN+^ genome-wide (Figure 3E, 3F). These strong, megadomain-specific biases were not mirrored in the fraction of 324 unique gene transcripts significantly down-regulated in *Setdb1*-deficient cortex (Figure 3F). Importantly, despite the spatial colocalization of the increased genic transcription in A_2_^NeuN+^ and the known ERV2 hotspots in B_2_^NeuN+^, we observed no fusion transcripts in any of the profiled RNA-seq samples (Table S13). Together, these findings strongly suggest that *Setdb1*-dependent epigenomic regulation of neuronal megadomains critically regulates A_2_^NeuN+^ and B_2_^NeuN+^ on a genome-wide scale (Figure 3G).

### Gliosis and genomic activation of microglia associated with IAP invasion of neuronal somata and processes

Analysis of the RNA-seq from *Setdb1* mutant as compared to control cortex revealed that the top 10 ranking gene ontology groups of up-regulated genes included regulators of ribosomal protein synthesis, the endoplasmic reticulum/endomembrane (ER/EM) stress response, ATP-dependent metabolism and the complement cascade (Figure S10, Table S14), potentially indicating a hypermetabolic state in response to an inflammatory stimulus. Strikingly, the cerebral cortex and striatum areas of *CamK-Cre^+^,Setdb1^2lox/2lox^* conditional mutant mice were affected by gliosis and exhibited upregulation of the astrocytic marker, glial fibrillary acid protein (GFAP) (Figure 4A, 4B). Similarly, Iba1 immunostaining marker revealed proliferative and reactive microglia in mutant hippocampus, although labeling of the overlying cortex was not significantly different from control (Figure 4A, 4C, 4D). Nonetheless, given the central role of microglia in brain inflammation, we conducted cell-type specific RNA-seq transcriptome profiling on immuno-panned microglia extracted from adult mutant forebrain of neuronal *Setdb1*-deficiency and controls (N=3/group). Expression of *Setdb1* transcript, including the loxP flanked exon 6 subjected to neuron-specific deletion in the mutant cortex, was completely preserved in the microglia from *CamK-Cre^+^,Setdb1^2lox/2lox^* mice (*Figure S11*). Furthermore, the overwhelming majority of retrotransposon transcripts, including the entire set of ERV2s, did not show elevated expression in microglia from mutant cortex (Table S15); these findings were expected given that *Setdb1* ablation is restricted to neurons in our conditional mouse model. We identified 840 (629 up, and 211 down) differentially regulated microglia-specific transcripts after neuronal *Setdb1* ablation (Figure 4E, Table S16). Among these were 629 up-regulated transcripts, for which gene ontology analyses indicated robust activation of interferon and cytokine signaling pathways and blood vessel formation including many genes associated with autoimmune and neuroinflammatory disease (Figure 4E, Table S16) (*21*). In contrast, no pathway enrichment was observed for the group of 211 microglial genes downregulated in mutant cortex. Next, given that the microglial genomic response to inflammatory stimuli also involves widespread changes in chromatin accessibility (*22*), we profiled open chromatin landscapes on a genome-wide scale using Assay for Transposase Accessible Chromatin (ATAC-seq) on CD11b-immunopanned microglia from mutant and control forebrain (N=3/group) (Figure 4F, Table S17). We identified 4154 open chromatin regions differentially regulated between two groups. Strikingly, microglial open chromatin region (OCR) upregulated after neuronal Setdb1 ablation showed the strongest enrichment (p<10^−43^) for binding motifs of Signal transducer and activator of transcription 2 (*Stat2*), a regulator of gene expression highly sensitive to activation by interferon and antiviral response pathways in brain and other tissues (*23, 24*). Furthermore, nuclear factor κB (NF-κB), a key factor in mediating the microglial genomic response to neuronal injury and inflammation (*25*), was among the top 5 ranking motifs in OCR upregulated in microglia from cortex with neuronal Setdb1-deficiency (Figure 4F). These findings, taken together, suggest that neuron-specific epigenomic un-silencing of ERV retrotransposons could trigger an inflammatory response with astrocytosis and microglial activation.

**Figure 4:**
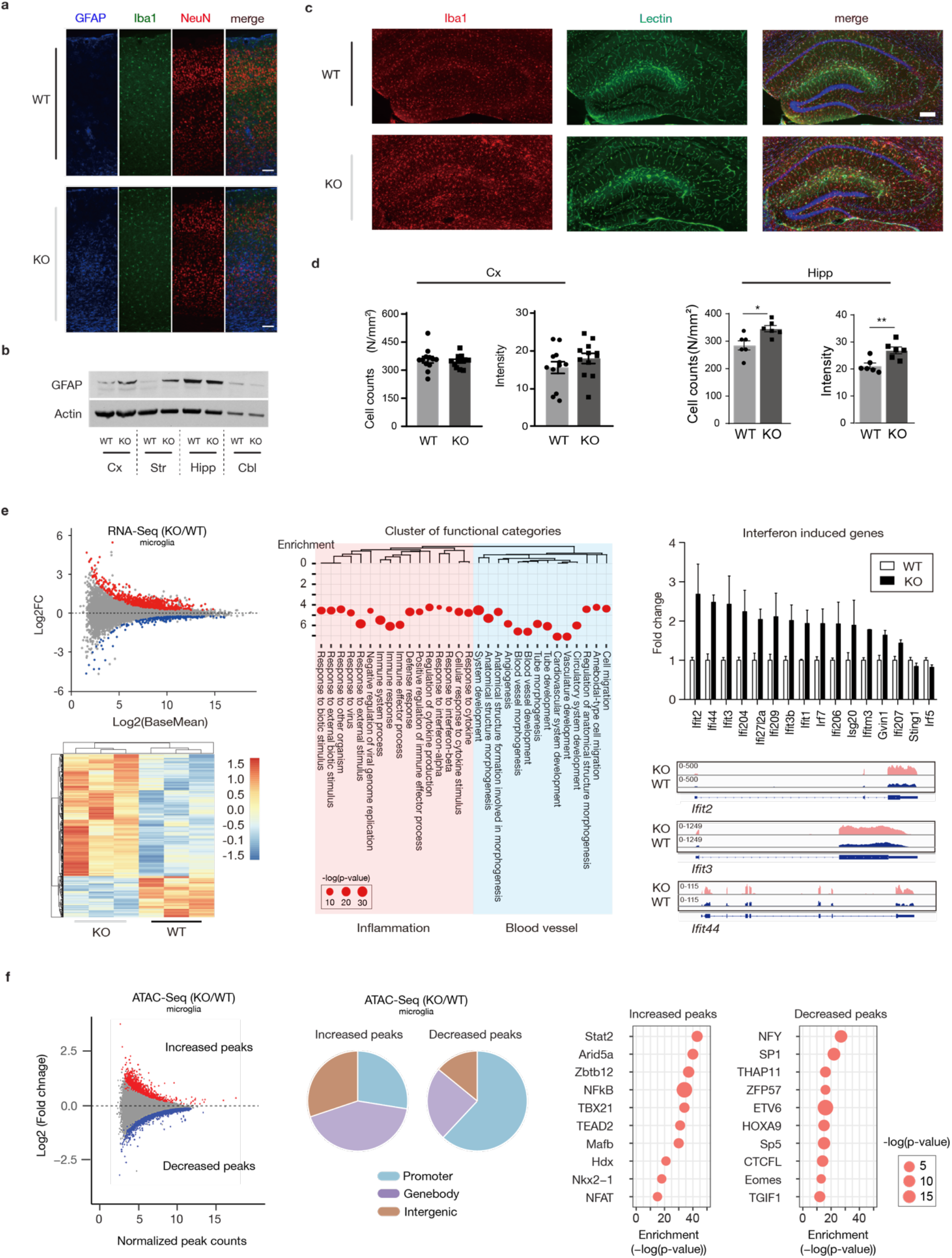
(A) Representative images show immunofluorescent signal for GFAP (blue), Iba-1(green) and NeuN (red) in cortex, from wildtype (WT) and *Setdb1* CK-cKO (KO) mice. Scale bar: 100 um. (B) Western blotting using the anti-GFAP antibody across several brain regions in WT and KO, as indicated. Cx, cortex; Str, striatum; Hipp, hippocampus; Cbl, cerebellum. b-actin included as a loading control. (C) Representative images show immunofluorescent signal for Iba-1 (red) and lectin (green) in hippocampus from wildtype (WT) and *Setdb1* CK-cKO (KO) mice. Cell nuclei counterstained with DAPI (blue). Scale bar: 200 um. (D) Bar graphs show quantifications of microglia cell number (left) and immunofluorescent intensity (right) in Cx (left) and Hipp (right) from WT and KO. Unpaired t-test. N=6 mice per group, *p < 0.05, **p < 0.01. (E) (Left) Top: MA-plot shows correlation between gene expression fold change (KO/WT) and abundance. Red, P<0.05, log2FC>0; blue, P<0.05, Log2FC<0; grey, P>0.05. Bottom: Clustered heatmap shows differential expression genes (P<0.05) in purified microglia from *Setdb1* wildtype (WT) and knockout (KO) forebrain (n=3/group). (Middle) ShinyGO analysis show functional enrichment in differential expression genes (KO/WT), FDR < 0.05, Log2FC > 0.5. (Right) Expression of significant interferon-induced genes with normalized FPKM in WT (white bar) and KO (black bar). (F) (Left) MA-plot show correlation between fold change (KO/WT) and base mean signal of ATAC-seq peaks in purified microglia from *Setdb1* wildtype (WT) and knockout (KO) forebrain. Red, P<0.05, Log2FC>0; blue, P<0.05, Log2FC<0; grey, P>0.05. (Middle) Genome annotation of up- and down-regulated significant peaks in KO as compared to WT. Blue, promoter; purple, genebody; brown, intergenetic. (Right) Homer de novo motif prediction for differentially regulated peaks. Notice inflammation related transcription factors including Stat2, Arid5a, and NFkB significantly enriched for up-regulated ATAC-seq peaks in KO. Size of the bubble (number in the key) indicates percentage of target sequence with predicated motif.

Studies in mice with severe immunodeficiencies and in genetically engineered cell lines suggest that the un-silencing of ERVs triggers an immune response primarily via RNA-sensing associated with the mitochondrial antiviral signaling protein (MAVS) and the Stimulator of Interferon Genes (STING) signaling pathways (*21, 26*). We observed that in the absence of Setdb1, there was increased transcription specifically of ERV2s in the setting of decreased H3K9me3 (Figure 5A); increased transcription of IAP-gag in cortical neurons was also visualized by RNA FISH (Figure 5B). To assess the viral burden in our mouse model, we next monitored protein expression and found that mutant, but not control cortex showed robust neuronal expression of the IAP Gag protein critical for retroviral assembly (Figure 5C, 5D). Electron microscopy confirmed dramatically increased numbers of mutant neurons with the presence of immature and mature IAP provirus in close proximity to cisterna-like membranous spaces compared to controls (Student’s two-sided t-test, p=3.69×10^−5^). In addition, counts of IAP particles (within the subset of IAP+ neurons in both genotypes) were much higher in mutants compared to controls (Wilcoxon sum-rank testing (p=0.008)) (Figure 5E). Strikingly, a subset of neurons showed dramatic proviral proliferation with encroachment into neuronal somata, dendrites and even spines in a subset of cortical neurons (Figure 5F). We conclude that the rewiring and epigenomic reorganization of ERV2-enriched neuronal megadomain sequences upon *Setdb1* ablation results in IAP retrotransposon escape from silencing. In addition to the robust increase in IAP transcript and protein levels, we observed proviral assembly and potentially dramatic accumulation of such particles in susceptible neurons resulting in viral hijacking of endomembrane systems, an ER/EM stress response, and ultimately neuroinflammation.

**Figure 5:**
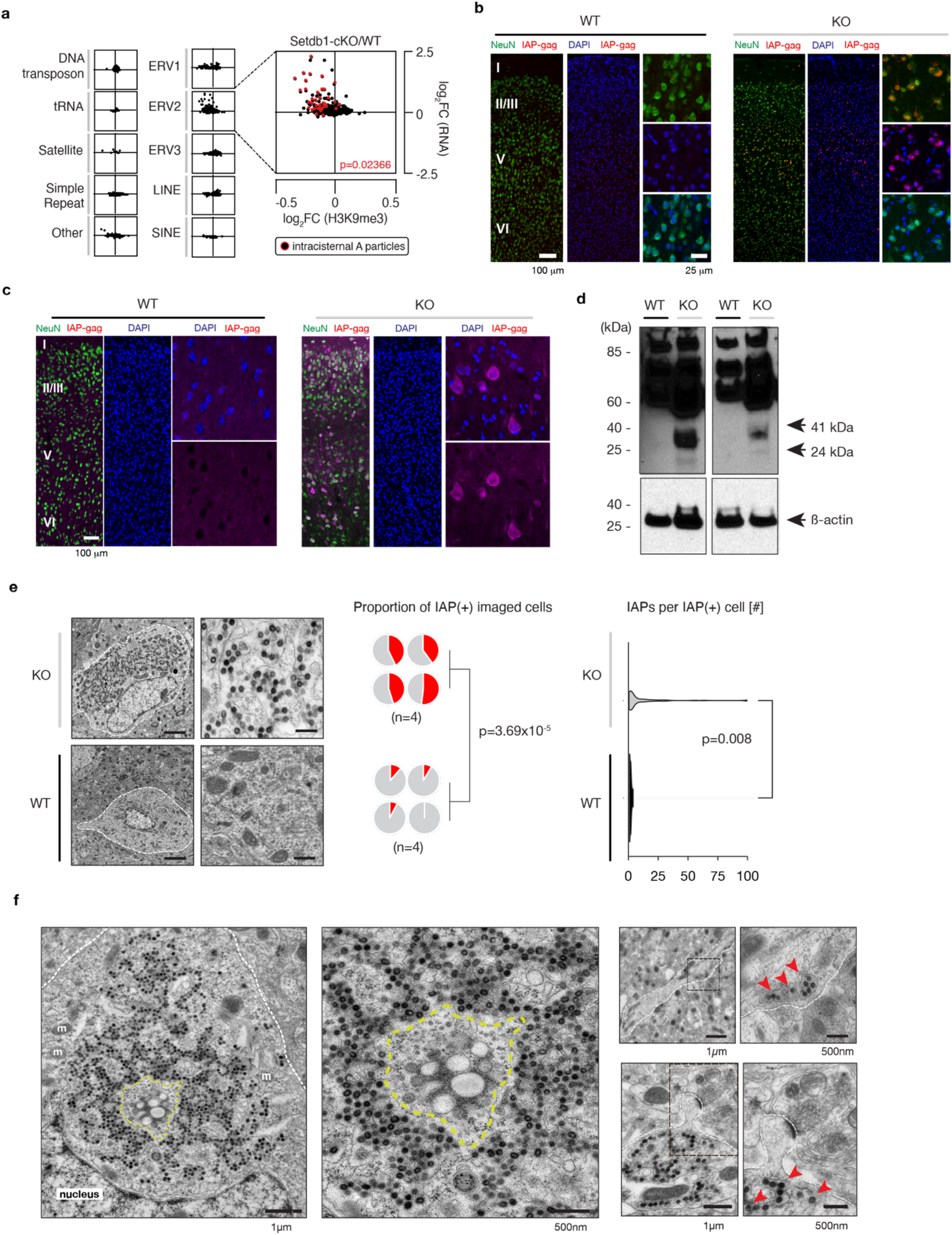
(A) Scatterplots of log2FoldChange (H3K9me3 KO vs. WT, x-axis) vs. log2FoldChange (RNA KO vs. WT, y-axis) for each of ten repetitive element categories (Repbase). The intracisternal A particles (IAPs) are labeled in red and overlaid on the ERV2 plot; IAPEzi as indicated. The significance of the regression is included (p=0.02366). (B) RNA FISH signal in coronal sections (30 um) across multiple cortical layers in KO vs. WT. Counterstained with NeuN (green) and DAPI (blue) (C) Representative immunofluorescence of WT (left) and KO (right) cortical cross sections stained for IAP-gag (pink) and NeuN (green), and DAPI counterstain. Scale bar, 50 μm. Cortical layers, Roman numerals I to VI. (D) Western blotting using the anti-IAP-gag antibody for full-length Gag (73kDa) and cleaved, processed mature forms (41kDa and 24kDa) in WT and KO mouse cortical tissue. b-actin included as loading control. (E) (Left) Cortical layers II/III electron microscopy (EM) images of KO (top) and WT (bottom) neurons imaged at 15 and 30k resolution. (Middle) Quantification of blinded counts of randomly imaged neurons (n=50 or 35) with >1 IAP particle present (IAP+ neurons) (n=4mice /genotype. (genotype [0 IAPs,1+ IAPs]: WT1 [31,4]; WT2 [32,3]; WT3 [47,4]; WT4 [51,0]; KO1 [20,15]; KO2 [21,14]; KO3 [28, 23]; KO4 [26,28]), Student’s two-sided t-test (p=3.69×10-5). (Right) Number of IAPs imaged within the IAP+ neurons, Wilcoxon sum-rank testing (p=0.008). (F) (Left) Sample neuron at low- and high-resolution with intracisternal proviral packaging and assembly hub marked with yellow dashed line; m, mitochondria; nucleus, cell nucleus. (Right, top) neuronal dendrites at 15k, 30k resolution. (Right, bottom) Low- and high-resolution images of IAPs at synaptic spine. Red arrows mark individual proviral particles. Scale bars as indicated.

## Discussion

We report that megadomain organization in the adult brain involves a unique, neuron-specific signature, including a B-type subcompartment encompassing 104Mb of neuronal chromatin (B_2_^NeuN+^) composed of comparatively ‘small’ (1-12Mb) heterochromatic ‘islands’ engulfed by A_2_^Neuron^ and other A-compartment-associated megadomains. Comparison of cell-type specific *SPRET/EiJ* vs. *C57Bl6/J* Hi-C maps, including mice with neuronal *Kmt1e/Setdb1* ablation, strongly points to epigenomic regulation of (germline-fixed) ERV2 transposable element sequences as a major driver for the B_2_^NeuN+^-to-B_2_^NeuN+^ and B_2_^NeuN+^-to-A_2_^NeuN+^ contact patterning.

Importantly, despite widespread rewiring and disruption of compartment-specific chromosomal patterning, *Setdb1*-deficient neurons only show few changes in the topologically-associating domain (TAD) landscape, with the notable exception of the *clustered Protocadherin* locus and a few additional genomic sites (*27*). This is consistent with the general notion that phase separation, as a molecular force, shapes compartments, as opposed to actively-driven loop extrusion presenting as TADs in ensemble Hi-C data, largely reflecting non- overlapping chromatin regulatory mechanisms (*28*). Thus, the strong interconnectivity of B_2_^NeuN+^ megadomains, and their regulatory effects on the surrounding A_2_^NeuN+^ subcompartment, could depend on ‘bridges’ of heterochromatin-associated protein 1 (HP1) bound to H3K9me2/3-tagged nucleosomes (*29*) and additional phase separation-promoting mechanisms (*30*) that, according to the present study, critically involve *Kmt1e/Setdb1*.

Our study strongly suggests that the reorganization of higher order chromatin in adult cortical neurons tracks the dramatic expansion of IAPs and other ERV2 transposable elements in *Mus musculus-*derived inbred lines, as compared to the wild-derived SPRET/Eij inbred strain harboring the 1.5 million year evolutionarilyy divergent *Mus spretus* genome (*18*). Previous studies linked a small subset of these newly inserted ERV sequences to strain-specific differences in gene expression, pathogenic mutations and neomorphisms (*31, 32*). However, our findings point to much broader cell-type specific adaptations in genome organization and function, including a partial remodeling of the chromosomal interaction map in mature cortical neurons. The importance of proper heterochromatic organization in neurons is underscored by the massive proliferation of IAP particles in a subset of the susceptible *Kmt1e/Setdb1* mutant neurons, in conjunction with high grade gliosis and reorganization of the neuronal transcriptome, reflective of an intracellular ER-driven stress response. In addition, secondary changes in gene expression programs in microglia were observed in response to the increased neuronal retrotransposon load. We note that, according to the cell-type specific single DNA sequencing of the present study, *Setdb1*-deficient neurons do not show a detectable increase in IAP/ERV2 *de novo* insertions, suggesting the dramatic observed ‘neuronal hijacking’ by the IAP proviral machinery is primarily driven by un-silencing of preexisting retrotransposon copies as opposed to excessive somatic retrotransposition events.

The findings reported are also of interest from the viewpoint of the potential neurotoxic effects of ERV2-like transcripts and proteins in human and invertebrate neurons, including potential links to tau- and TDP-43 associated neurodegenerative disease (*33–37*), and a recent report on increased HERV activation in the adolescent non-human primate brain exposed to maternal immune activation during prenatal development (*38*). Interestingly, we observed enrichment for Human ERV2-like (HERV-K) repeat sequences in inter-autosomal Hi- C contact maps from human neurons, an effect driven primarily by loci with strong synteny with murine B_2_ megadomains (Figure S12, S13), strongly suggesting that compartment-specific enrichment of ERV2 sequences occurs in parallel across different mammalian lineages. Therefore, it is plausible that cell-type specific 3D genome organization in the mammalian brain includes a type of heterochromatic organization adaptive to species- and strain-specific reconfiguration of the ERV retrotransposon landscape, and highly relevant for brain function in health and disease.

## ACKNOWLEDGMENTS

The authors thank all members of the Akbarian lab for constructive comments and discussions; E. Xia, C. Rosenbluh, and A. Chess for generous access to sequencing equipment; L. Winterkorn and staff at the New York Genome Center for logistical support; M. Smith and D. Sachs for guidance on the PacBio SMRT-Seq protocol; and L. Mulder for his insights into human retrovirology. Nuclei sorting was performed at the Flow Cytometry CoRE at the Icahn School of Medicine at Mount Sinai. Microscopy preparation and imaging was performed at the Microscopy CoRE at the Icahn School of Medicine at Mount Sinai. RNA-Seq library preparation and next-generation sequencing was performed at the Genomics CoRE at the Icahn School of Medicine at Mount Sinai. We thank Dr. B.R. Cullen (Duke University) for the anti-IAP antibody gift.

## Author contributions

S.C., B.J., B.K., Y.D., Y.Z., L.K.B., P.R., C.J.P., E.A-P., and M.I. performed experiments; S.C. and S.A. conceived and designed experiments; S.C. performed statistical analyses; S.C., S.E-G., Y-H.E.L., Y.D., M.E., and L.S. performed bioinformatics and genomic analyses; Y.J. and S.A. supervised the research; S.C. and S.A. wrote the paper, with contributions from all co-authors.

## Data and materials accessibility

Scripts used for bioinformatics analyses will be made available at https://github.com/sandchand5/hic-neuron-erv upon acceptance for publication. Sequencing data for genome-scale analysis (Hi-C, ChiP-Seq, RNA-Seq, PacBio SMRT-Seq) have been deposited in NCBI’s Gene Expression Omnibus and will be made accessible prior to publication. All other data included in the publication and are available from the authors upon request.

## Supplementary Materials

**This PDF file includes:**

Materials and Methods

<u>Figures S1 to S13</u>

Figure S1 NeuN+ Hi-C clusters

Figure S2 NeuN− Hi-C clusters

Figure S3 Trans interactions by subcompartment

Figure S4 Trans interactions among B2 chromosomal megadomains

Figure S5 Gene clusters in B2

Figure S6 CTCF binding motif enrichment

Figure S7 ERV2 hotspots interact least frequently in SPRET/EiJ NeuN+

Figure S8 PacBio SMRT long-read sequencing of IAPEzi *de novo* integration sites

Figure S9 Hi-C circos plots from NeuN+ of *Camk-Cre^+^, Setdb1^2lox/2lox^* mutant mice (adult cortex)

Figure S10 GOs for elevated transcripts in *Camk-Cre^+^, Setdb1^2lox/2lox^* mutant vs. control adult cortex, divided by subcompartment

Figure S11 Microglia RNA-Seq and ATAC-Seq

Figure S12 Mouse/human subcompartment comparisons

Figure S13 Inter-chromosomal contacts in human neurons are enriched with (H)ERV retroelements

**Other Supplementary Materials related to this manuscript include the following and will be provided as an .xlsx file upon request:**

<u>Tables S1 to S17</u>

Table S1 Hi-C QC metrics

Table S2 NeuN+ subcompartment genomic coordinates (GRCm38/mm10)

Table S3 NeuN− subcompartment genomic coordinates (GRCm38/mm10)

Table S4 ChIP-Seq metrics

Table S5 NeuN+ trans interactions

Table S6 NeuN− trans interactions

Table S7 Repeat Sequence Classification

Table S8 PacBio SMRT long-read sequencing QC metrics

Table S9 PacBio full-length coordinates (de novo)

Table S10 Differential trans interactions Setdb1-deficient and control neurons with wildtype Setdb1 levels (padj < 0.05)

Table S11 Differential cis interactions Setdb1-deficient and control neurons with wildtype Setdb1 levels (padj < 0.05)

Table S12 RNA-Seq metrics

Table S13 Discordant + chimeric RNA-Seq

Table S14 Genomic RNAs

Table S15 Differential DNA repeats of RNA-Seq from Setdb1-deficien and control microglia (p < 0.05)

Table S16 Differential genes list of RNA-Seq from Setdb1-deficient and control microglia (p < 0.05)

Table S17 Differential peaks list of ATAC-seq from Setdb1-deficient and control microglia (p < 0.05)

### Mouse handling

All mouse work detailed below was approved by the Institutional Animal Care and Use Committee of the Icahn School of Medicine at Mount Sinai. Mice were held under specific pathogen-free conditions with food and water being supplied *ad libitum* in an animal facility with a reversed 12h light-dark cycle (lights off at 7:00 AM) under constant conditions (21 ± 1C, 60% humidity). All mice were group-housed (2-5 mice per cage). For use in our experiments, mice were reared into adulthood (≥3 months) and were sacrificed by decapitation following isoflurane anesthetization according to IACUC guidelines. Following brain removal, cortical dissections were performed on ice aided by a dissection light microscope.

### Cell-type specific Hi-C

#### Fluorescence activated nuclear sorting

Cortical tissue was prepped for fluorescence activated nuclear sorting (FANS) as follows. Briefly, the dissected tissue was homogenized in a hypotonic lysis solution and fixed in 1% formaldehyde for 10 minutes at room temperature. The cross-linking reaction was quenched with 125 mM glycine. The nuclei were then purified by centrifugation at 4000xg and resuspended in a 1:1 solution of the hypotonic lysis solution and a 1.8M sucrose solution prior to re-centrifugation at 4000xg to isolate out cortical nuclei. The pellet was then resuspended in Dulbecco’s phosphate buffered saline (DPBS) containing 0.1% BSA and 1:1000 anti-NeuN antibody (clone A60, Alexa Fluor 488 conjugated; EMD Millipore Corp., MAB377X). Samples were incubated for 45 minutes while rotation and protected from light at 4C. DAPI (Invitrogen) was added immediately before sorting to label all nuclei. Sorting was performed at the Flow Cytometry CoRE at the Icahn School of Medicine at Mount Sinai. Nuclei were collected as NeuN+ and NeuN− populations following serial gating and pelleted for downstream experimental processing.

##### in situ Hi-C

Nuclei were digested with 100U MboI, and the restriction fragment ends were labeled using biotinylated nucleotides and re-ligated. After reversal of the cross-linking, ligated DNA was purified and sheared to a length of ∼400bp by sonication, and the biotin-tagged ligation junctions were subsequently pulled down with streptavidin beads (Invitrogen, Dynabeads MyOne Streptavidin T1, Catalog No. 65602) and prepared into libraries for next-generation Illumina sequencing (HiSeq 2500). For the SPRET/Eij and matched C57/BL6J studies, the Arima Hi-C kit (Arima Genomics, San Diego) was used according to the manufacturer’s instructions.

### Hi-C bioinformatics

#### HiC-Pro

The ‘pre-truncation method’ using HiC-Pro(v2.9) was used for quality control purposes. Libraries were mapped to the *Mus musculus* reference genome (GRCm38.p5_M13) using bowtie2.2. Artifacts and other common statistics are available as Supplementary Information. HiC-Pro results were piped to Juicer to produce .hic format files using the parser tool ‘*hicpro2juicebox*’. These files, which contain compressed contact matrices at varying resolutions, were then processed into final matrix-balanced normalized contact maps with Juicebox.

#### HOMER

Each read of the paired-end libraries was aligned independently using bwa-mem(v0.7.15), permitting split-mapping to the GRCm38.p5_M13 reference annotation. After mapping, forward and reverse reads were directly supplied to Homer(v4.8) for processing by first, merging the paired-end reads and later, filtering out self-ligation (spikes and continuous) artifacts. Files were normalized based on sequencing depth and distance between loci, creating a background model necessary for calculating significant pairwise interactions. Trans- interactions (1Mb resolution) were similarly calculated, but for significant interactions for loci >200Mbp apart (*-minDist* 200000000) to disregard intra-chromosomal interactions.

#### Subcompartment calling

We adapted a previously published k-means clustering algorithm(*7*) to identify subcompartments within our NeuN+ nuclei. In short, a 250kb autosomal resolution map was constructed from NeuN+ and NeuN− validPairs.hic files generated with HiC-Pro(v2.9). The matrices were normalized with matrix-balancing and processed as observed/expected matrices. Specifically, a subset of interchromosomal contact data was extracted using .hic dump and stitched together, with 250kb loci on odd-numbered chromosomes serially appearing as rows and 250kb loci on even-numbered chromosomes serially appearing as columns. Odd chromosomal loci were first clustered using the *kmeans* function in R, and even chromosomal loci were similarly clustered after transposing the stitched matrix. Several values for the clustering parameter, “k”, were tested, ranging from k=2 to k=10 to forcibly organize the genomic loci across these chromosomes into *k* number of clusters. For odd chromosomes, a cluster number of k=6 was determined as the best fit by visual inspection with the generated Hi-C raw read contact matrices; similarly, for even chromosomes, a cluster number of k=5 was determined as optimal. Within each of these cluster sets, clusters consisting of <5% of the genome were disregarded. This ultimately resulted four clusters for the odd and even chromosome sets, that were merged in order of size to represent four subcompartments for the entire genome.

#### Subcompartment characterization

To appropriately classify these subcompartments as active (‘A’) or inactive (‘B’), we leveraged publicly accessible data from the mouse ENCODE project (led by the Mouse ENCODE Consortium). The database is a curation of numerous epigenetic profiles, including transcription factor and polymerase occupancy (http://chromosome.sdsc.edu/mouse/download.html), DnaseI hypersensitivity, histone modifications, and RNA transcription. Strikingly, our results showed a clear dichotomy with respect to overlap with lamina-associated domains (*39*) wherein two subcompartments exhibited a majority overlap with LADs (‘B’), and two did not (‘A’). Euchromatin-associated epigenetic features, both active and facilitative (enhancers, CTCF, H3K4me3 *(ENCFF572BJO)*, H3K4me2 *(ENCFF647TKR)*, H3K9ac *(ENCFF540IHT)*, H3K4me1 *(ENCFF224LTF)*, H3K27ac *(ENCFF676TSV)*, RNAPolII, and H3K36me3 *(ENCFF999HGN)*), showed a near complete overlap with the ‘A’ subcompartments, while the smaller of the ‘B’ subcompartments showed moderate overlap with the repressive modification H3K9me3 *(ENCFF101VZW)*. Consequently, the derived subcompartments were delineated as ‘A’ (euchromatic) or ‘B’ (heterochromatic), and numbered in order of decreasing size, resulting in ‘A1’, ‘A2’, ‘B1’, and ‘B2’. Notably, blacklisted ENCODE coordinates (portions of the genome that consistently produce anomalous high signal/read counts) accounted for <0.01% of the genomic space encompassed within any of the determined subcompartments; this minimizes the possibility that the signal enrichments determined within these subcompartments are products of aberrant read mapping.

#### Hi-C *trans* and *cis* calculations

The aforementioned 250 kb autosomal, matrix-balanced resolution maps of observed/expected Hi-C interaction frequencies were used for downstream analyses; each locus along the axes of the matrix was then labeled according to its NeuN+ subcompartment identifier (A1, A2, B1, B2). For *trans*, Hi-C values from combinations of pairwise interactions involving loci of specific subcompartment designations on different chromosomes were extracted and populated into a vector; for *cis*, these values were extracted for loci on the same chromosome, excluding the diagonal.

#### SPRET/EiJ vs. C57/BL6J Hi-C comparison

Hi-C libraries were generated for sorted NeuN+ and NeuN− populations from SPRET/EiJ and C57/BL6J mice (age- and sex- matched) using the Hi-C Next Generation Sequencing Kit (Arima Genomics) according to the manufacturer’s protocol. Next, 250 kb autosomal resolution maps were constructed for using valid pairs generated from the HiC-Pro (v2.9) pipeline following alignment to the *mm10* reference genome. The matrices were normalized with matrix-balancing and processed as observed/expected matrices. Each locus along the axes of the matrix was then labeled according to its NeuN+ subcompartment identifier (A1, A2, B1, B2) and ERV2 count (as determined from UCSC Repeatmasker). A value of 88 was calculated to reflect a locus harboring the 99^th^ percentile of ERV2s as compared to the rest of the genome, and loci with >99^th^ percentile of ERV2s were retained for downstream analyses, generating a reduced Hi-C matrix of 101×101 loci for each sample. Samples were combined by cell type and strain, resulting in four final matrices representing SPRET/EiJ NeuN+, C57/BL6J NeuN+, SPRET/EiJ NeuN−, and C57/BL6J NeuN−. Statistical significance was performed using Wilcoxon sum-rank testing (paired, two-sided).

#### Differential trans- and cis- analysis

Raw *in situ* Hi-C matrices (20kb resolution) were binned into 1Mb segments genome-wide, and DESeq2 was used to determine windows of significant differential read scores in *trans* (p_adj_ < 0.05). Interaction anchors and targets were classified by subcompartment based on percentage of the bin that encompassed subcompartment loci; in the event of a tie, the assignment was made hierarchically in order of subcompartment size, smallest to largest. Expected values of interactions for subcompartment anchor-target combinations were determined from the relative sizes of the subcompartments of interests. Pearson’s residuals for differential were calculated by subtracting the residuals of decreased Hi-C interactions

### Chromatin immunoprecipitation studies

#### Native ChIP-seq (NChIP-seq)

10^6^ NeuN+ and NeuN− nuclei were pelleted after FANS, resuspended in 300µl of micrococcal nuclease digestion buffer (10mM Tris pH 7.5, 4mM MgCl, and 1mM Ca^2+^), and digested with 2 uL of MNase (0.2U/µl) for 5 minutes at 28C to obtain mononucleosomes. The reaction was quenched with 50mM EDTA pH 8. Chromatin was released from nuclei with the addition of hypotonic buffer (0.2mM EDTA pH 8, containing PMSF, DTT, and benzamidine). Chromatin was incubated with the appropriate antibodies (anti-H3K9me3 (Abcam, ab8898), anti- H3K27ac (Active Motif, #39133), or anti-H3K79me2 (Abcam, ab3594)) overnight at 4C. The DNA-protein-antibody complexes were captured by Protein A/G Magnetic Beads (Thermo Scientific 88803) by incubation at 4C for 2h. Magnetic beads were sequentially washed with low-salt buffer, high-salt buffer, and TE buffer. DNA was eluted from the beads, treated with RNase A, and digested with Proteinase K prior to phenol-chloroform extraction and ethanol precipitation. For library preparation, ChIP DNA was end-repaired (End-it DNA Repair kit; Epicentre) and A-tailed (Klenow Exo-minus; Epicentre). Adapters (Illumina) were ligated to the ChIP DNA (Fast-Link kit; Epicentre) and PCR amplified using the Illumina TruSeq ChIP Library Prep Kit. Libraries with the expected size (∼275bp) were submitted to the New York Genomics Center for next-generation Illumina sequencing (HiSeq 2500, 75bp, paired-end).

#### Bioinformatics

To perform these bioinformatics analyses, we first aligned the raw.fastq reads to the *Mus musculus* reference genome (mm10) using bowtie2. Only concordant reads were compressed into aligned .bam files then sorted and indexed using the *samtools/0.1.19* suite, and following PCR duplicate removal, were converted into .bed files for further processing.

#### Cell-type specific histone profiling

We performed diffReps(*40*) differential analysis at a 1kb resolution for each histone mark of interest at a statistical significance threshold of p<0.001, with NeuN+ histone modification profiles as our treatment group, and NeuN− histone modification profiles as our control group. Significantly called regions (spanning a length of one kilobase of genome or greater) were labeled as NeuN+ enriched or NeuN− enriched according to their log2FoldChange values.

#### Genome association testing

The *Mus musculus* mouse (mm10) reference genome was divided into 250kb bins using tileGenome() in the GenomicRanges R package. diffReps output files were overlapped with these bins using the sum(countOverlaps) function to retrieve the number of diffReps regions within each 250kb bin. The two considered variables for the Fisher’s testing were (1) the subcompartment identifier of the 250kb bin and (2) the classification of the 250kb bin as ‘high’ or ‘low’ as it pertained to epigenetic feature enrichment. The threshold distinguishing ‘high’ and ‘low’ was set as the 99^th^ percentile of diffReps loci (or repetitive elements) within a 250kb bin across the autosomal reference genome. Fisher’s testing was performed using the *exact2×2* R package. For association with Hi-C interaction changes, bins involved as anchors or targets in the interactions were grouped as the reference genome set. For RNA association testing, no percentile threshold was used for filtering ‘high’ and ‘low’; instead any bin with altered RNA (either increased or decreased, as indicated) was considered ‘high’, with bins harboring no altered RNA designated ‘low’.

### PacBio SMRT Sequencing

#### Library preparation

SMRTbell libraries were constructed from IAPEz-int xGen Lockdown probe-captured DNA for sequencing using a PacBio Sequel II System. In short, genomic DNA was isolated from FACS-sorted NeuN+ or NeuN− nuclei of adult mouse cerebral cortex using phenol chloroform extraction and was subsequently sheared to ∼10kb using g-Tube microcentrifuge tubes (*Covaris, 010145*). End-repair, A-tailing, adapter ligation (barcoding) and PCR amplification with the universal primer were performed according to protocol, and fragments were appropriately size-selected using AMPure SPRI Select beads. Samples were then pooled prior to hybridization with biotinylated xGen® Lockdown® designed against the consensus sequences (dfam.org) of IAPEzi or IAPEY3- int (control) (Integrated DNA Technologies). Streptavidin A1 beads were used to capture hybridized fragments, which were subsequently amplified prior to SMRTbell library preparation. Following AMPure purification and size selection, SMRTbell templates were annealed and bound to the barcoded libraries, which were submitted for PacBio Sequel II HiFi sequencing (Genewiz, 30h movies).

#### Bioinformatics

Bioinformatics processing was performed with SMRT Link v5.1.0, run with default parameters, unless otherwise indicated. Files were first demultiplexed with *lima* and circular consensus sequences (CCS) were generated with *ccs* for filtered sequences with matching paired-end adapters. CCS reads were then mapped to an IAPEzi consensus sequence (*dfam.org*) using *pbalign*. Passing reads were subsequently mapped to the mouse reference genes using bwa split mapping and filtered based on mapping score (=60). Reads were then assigned as autosomal IAPEzi or autosomal non-IAPEzi IAP using the GenomicRanges subsetByOverlaps() function. The other reads, those not mapping to IAPs denoted in the reference genome, were then blasted against the IAPEzi consensus sequence and IAP masked reference genome in parallel; reads with higher bit scores for the IAPEzi blast than the reference genome were defined as *de novo*. Full-length *de novo* insertions were defined as fragments representing >5833bp (0.9*6481).

### Cortical Transcriptomics

#### RNA-seq

Total RNA was first extracted from the prefrontal mouse cortices (n=12, 6WT/6KO), and prepared with RNeasy Lipid Tissue Mini kit (with on-column DNase1 treatment). The quantity and quality of RNA was checked using a bioanalyzer (Agilent RNA 6000 Nano Kit). 1.5 µg of total RNA from each sample was submitted to the Genomics CoRE Facility at Mount Sinai for RNA-seq library generation with RiboZero treatment, and were subsequently sequenced on the Illumina HiSeq 2500, 100bp, paired end.

#### Bioinformatics

Read pairs were aligned to the *Mus musculus* mouse reference genome (mm10) Tophat2 short-read aligner. Reads were counted using HTSeq against the Gencode vM4 Mouse annotation. Genes were filtered based on the criteria that all replicates in either condition must have at least 5 reads per gene. On the resulting filtered transcript, a pairwise differential analysis between Setdb1 conditional mutant vs control cortex was performed using the voomlimma R package8,9 which converts counts into precision weighted log2 counts per million and determines differentially expressed genes using a linear model. Significantly differentially expressed genes were identified using a cutoff of Benjamini-Hochberg adjusted p-value less than 0.05. Discordant pairs were extracted from aligned .bam files using *samtools* with the following command: samtools view –b –F 2 and reads identifiable with specific flags were quantified and categorized.

#### Subcompartment neighbor testing

Subcompartment bins (250kb) were first determined and reduced using GenomicRanges into contiguous segments of similar identity. The percent of all boundaries by different combinations of subcompartment boundaries were calculated as the observed values. For expected values, genomes were reconstructed using the 250kb blocks and reduced as above. The percent of boundaries by different combinations of subcompartment boundaries were calculated x 100 permutations using regionER resampleRegions (mean + standard deviation).

#### GO Enrichment

Gene ontologies for gene sets of interest were determined using the ClueGO (v2.5.1) application through Cytoscape (v3.6.1). All available ontologies/pathways (CORUM, Chromosomal-Location, GO, INTERPRO, KEGG, REACTOME, WikiPathways) were considered, and the chosen network specificity was ‘medium’, displaying only pathways with pV<0.05. The statistical testing performed was an enrichment (right-sided hypergeometric test) analysis with Benjamini-Hochberg pV correction. All other parameters were left as default parameters, including GO Term Grouping.

### Microglial Studies

#### Microglia Isolation

Single-cell suspension from adult brain tissues was prepared using Miltenyi’s Adult Brain Dissociation Kit (Miltenyi Biotec, 130-107-677) and Debris Removal Solution Kit (Miltenyi Biotec, 130-109-398) according to manufacture instruction with minor modifications. In brief, total brain tissues, except olfactory bulb and cerebellum, were collected quickly and washed with ice cold 1x HBSS. After chopped into small pieces using sharp blade, the brain tissues were transferred into the C-tube containing 1950ul of Enzyme mix 1 and 30ul of Enzyme mix 2 from the kit and incubated on Miltenyi’s gentle MACS Octo Dissociator with Heaters using program 37°C _ABDK_1 for 30 minutes. Afterwards, the digested tissue homogenate gently went through fire polished glass pippette 10 times, and then passed through a 70um cell strainer. Cells were collected via centrifuged at 300g for 10 minutes at 4 °C and resuspended in ice cold 1x HBSS. Debris Removal Solution was then added and overlayed with ice cold 1x HBSS. After centrifugation, three phases in the tube were clearly visualized, from top to bottom: 1x HBSS solution, debris and myelin layer, and single-cell suspension. Discard the top two phases completely, wash the cells in 1x HBSS, centrifuge to collect the cells, and then resuspended the cell pellet in 1 ml of 1x HBSS containing 1% fetal bovine serum. In order to minimize the unwanted microglia activation, single cell suspension was incubated in Fc receptor blocking Reagent (Miltenyi Biotec, 130-092- 575) for 10 min, and then the cell suspension was incubated with CD11b-microbeads (Miltenyi Biotec, 130- 093-634) for 10 min at 4 °C in the dark, followed by positive selection with LS separation column (Miltenyi Biotec, 130-042-401). Flow cytometry analysis was performed to check the cell purity. The three different groups of cells (no enrichment, target and non-target cell population) were incubated in CD11b-FITC Monoclonal Antibody (M1/70) (eBioscience, 11-0112-81) at 1:2000 dilution for 30 min at 4 °C in the dark. Stained cells were examined using the Beckman flow cytometer.

#### Immunostaining

The coronal sections (30um) were washed with 1x PBS containing 0.5% Triton X-100 for 10 min, and then blocked with 1x PBS containing 0.5% Triton X-100 and 5% goat serum, followed by primary antibodies incubation for anti-Iba1 (Abcam, ab178846), anti-GFAP (Abcam, ab4674), anti-NeuN−Alex555 (Millipore, MAB377X), Tomato-lectin (Vector, DL-1174,) at 4°C overnight. Sections were incubated with fluorescence-labeled secondary antibodies at room temperature for 2 h in the dark. Images were captured with a fluorescence microscope (Olympus VS120). Analysis was performed using NIH Image J software.

#### RNA-seq

Total RNA was extracted by using Direct-zol RNA MicroPrep (Zymo research, R2060). Library was prepared using QIAseq FastSelect RNA Removal Kit (Qiagen, 333180-24) and QIAseq Stranded Total RNA Lib Kit (Qiagen, 180743) following manufacturer’s instruction. In brief, 200-500ng total RNA was used for each reaction. 1ul rRNA removal reagent was added into 28ul of RNA sample, plus 5ul 5 x RT Buffer, followed by incubation for 3min at 95°C, and then went through stepwise annealing using PCR programing. After fragmentation and rRNA removal, reverse transcription, second-strand synthesis, end-repair, A-addition, and strand-specific ligation with selected adapter (1:25 dilution) were performed, followed by CleanStart library amplification. Library DNA was then purified and size selected for around 500bp fragments, and checked with Qubit and Agilent 4200 Tapestation.

#### ATAC-seq

Cells were collected and nuclei were extracted by douncing directly in lysis buffer (0.32M sucrose, 5mM CaCl2, 3mM Mg(Ace)2, 0.1mM EDTA, 10mM Tris-HCl pH=8, 0.1% NP-40. 50,000 nuclei were used for each reaction. Library was prepared using TruePrep DNA Library Prep Kit V2 for Illumina (Vazyme, TD501-02) following manufacturer’s instruction with minor modifications. In brief, nuclei were treated with Tn5 at 37°C for 30 min, and then DNA was purified with MinElute Gel Extraction Kit (Qiagen, 28604), and elute in 25ul of EB buffer. PCR was then setup by the mixture of 20ul DNA sample, 10ul TAB, 5ul PPM, and selected adaptors (5ul N5XX and 5ul N7XX) (TD202), followed by PCR amplification. Library DNA was purified by using SPRIselect beads (Beckman, B23318) for 1:0.55 followed by 1:1.15 two steps size selection, and then checked with Qubit and Agilent 4200 Tapestation.

### Microglial Data Analysis

#### RNA-seq

Sequencing was performed on Illumina XTen (PE150). Before analyzed the data, FastQC was first used for quality control analysis, and Trim-galore was used to remove low-quality reads. Paired-end clean data was aligned to reference genome (M. musculus, UCSC mm10) using Tophat2 v2.1.1. Samtools v1.9 was used to sort and build the alignment files index. FeatureCounts v1.6.3 from subread package was used to get gene expression level counts by determining the number of reads mapped to gene exons, and differential analysis was generated by DESeq2. Significant genes (log_2_FoldChange > 0.5, Padj < 0.05) were extracted for Gene Ontology Enrichment Analysis by using ShinyGO v0.61 (http://bioinformatics.sdstate.edu/go/). Deeptools was used to generate normalized bigwig and visualized on IGV.

#### ATAC-seq

Sequencing, FastQC, and data cleaning were performed as described above. Paired-end clean data was then aligned to reference genome (M. musculus, UCSC mm10) using Bowtie2 v2.3.4.3. Samtools v1.9 was used to sort and build the alignment files index after removing duplications. MACS2 was used for peak calling (--shift - 100 --extsize 200 --nomodel). Diffbind was used to generated differential change peaks. R packages “ChIPseeker”, “clusterProfiler” “org.Mm.eg.db” and “TxDb.Mmusculus.UCSC.mm10.knownGene” were used for peak annotation. Significant differential peaks that are located on gene promoters were extracted to generate Gene Ontology Enrichment Analysis by ShinyGO v0.61 (http://bioinformatics.sdstate.edu/go/). Deeptools was used to generate normalized bigwig and visualized on IGV.

### RNAScope in situ hybridization of IAP-gag probes

Coronal mouse brain slices (10 um) were collected using a freezing microtome and mounted onto Superfrost plus slides (Fisher). The slides were processed per the RNAscope Multiplex Fluorescent v2protocol (Advanced Cell Diagnostics). Briefly, the tissue sections were treated with protease, hydrogen peroxide, and incubated with either sense probes for DNA FISH, or anti-sense probes for RNA FISH, targeting the IAP-gag sequence for 2 hours at 40°C. Amplifier sequences were polymerized to the probes and treated with Opal 570 dye. The slides were incubated in NeuN−AlexaFluor488 antibody (1:200 in 1xPBS; MAB377X; EMD Millipore) for 2 hours at room temperature and cover-slipped with DAPI Fluoromount (Southern Biotech). Imaging was performed on a Zeiss LSM780 confocal microscope.

### Protein expression

#### Immunohistochemistry and Western blotting

Coronal sections from perfusion-fixed (by phosphate-buffered 4% paraformaldehyde) adult mouse brains were processed for anti-GFAP immunoactivity and detected with diaminobenzidine (DAB) using the ABC kit (VectorLabs) according to the manufacturer’s protocol. For immunoblotting, protein was extracted from homogenized adult mouse cortical tissue and 100 microgram total protein was loaded in each lane. The membrane was blotted with rabbit anti-IAP antibody (a gift from Dr. Bryan R. Cullen, Duke University, 1:10000 dilution) and probed with goat anti-rabbit HRP (1:5000 dilution) for one hour at room temperature prior to detection.

For IAP immunohistochemistry, adult conditional *Setdb1* mutant and control mice - approximately 6 months old - were anesthetized with a terminal intraperitoneal injection of a ketamine/xylazine mixture (IP: 200 and 30 mg/kg, respectively). Transcardial perfusion was performed with 100ml of 10% sucrose followed by 200ml of 4% paraformaldehyde in PBS. Brains were removed and placed in 4% formaldehyde overnight at 4°C, followed by incubation in 30% sucrose until isotonic. After embedding in OCT compound (Tissue-Tek), the brains were cut on a freezing microtome (Leica SM2010 R) into 30 µm coronal sections and placed in 1x PBS. Staining for IAP protein was performed as follows: coronal sections containing prefrontal cortex were blocked and permeabilized with 10% BSA and 0.05% Triton X-100 in 1x PBS for 1 hr at room temperature, followed by incubation with the rabbit anti-IAP antibody (see paragraph above) (1:100 in 1x PBS; and 0.01% Triton X-100 overnight at room temperature. The sections were washed for 5 minutes in PBS followed by incubation in anti-rabbit Alexa Fluor 647 secondary antibody for 1 hour at room temperature. Sections were washed briefly in PBS before being mounted on Superfrost Plus slides (Fisherbrand) with DAPI Fluoromount-G media (SouthernBiotech). Imaging was done using a Zeiss CLSM780 upright microscope.

### Electron Microscopy

Adult mice (N=4 conditional *Setdb1* mutants and N=4 control animals) were anesthetized and perfused using a peristaltic pump at a flow rate of 35 mls/min with 1% paraformaldehyde/phosphate buffered saline (PBS), pH 7.2, and immediately followed with 2% paraformaldehyde and 2% glutaraldehyde/PBS, pH 7.2 at the same flow rate for an additional 10-12 minutes. The animal’s brain was removed, and placed in immersion fixation (same as above) to be post-fixed for a minimum of one week at 4 degrees C. Fixed brains were sectioned using a Leica VT1000S vibratome (Leica Biosystems Inc., Buffalo Grove, IL) and coronal slices (400 ums) containing the frontal cortex were removed and embedded in EPON resin (Electron microscopy Sciences [EMS], Hatfield, PA). Briefly, sections were rinsed in buffer, fixed with 1% osmium tetroxide followed with 2% uranyl acetate, dehydrated through ascending ethanol series and infiltrated with EPON resin (EMS). Sections were transferred to beem capsules, and heat polymerized at 60 degrees C for 48 - 72 hours. Semithin sections (0.5 and 1 um) were obtained using a Leica UC7 ultramicrotome, counterstained with 1% Toluidine Blue, cover slipped and viewed under a light microscope to identify and secure the layers of interest (L11 – L1V). Ultra-thin sections (80nms) were collected on copper 300 mesh grids (EMS) using a Coat-Quick adhesive pen (EMS), and serial sections were collected on carbon-coated slot grids (EMS). Sections were counter-stained with 1% uranyl acetate followed with lead citrate and imaged on a Hitachi 7000 electron microscope (Hitachi High-Technologies, Tokyo, Japan) using an advantage CCD camera (Advanced Microscopy Techniques, Danvers, MA). Images were adjusted for brightness and contrast using Adobe Photoshop 11.0.

---

**Figure S1.**
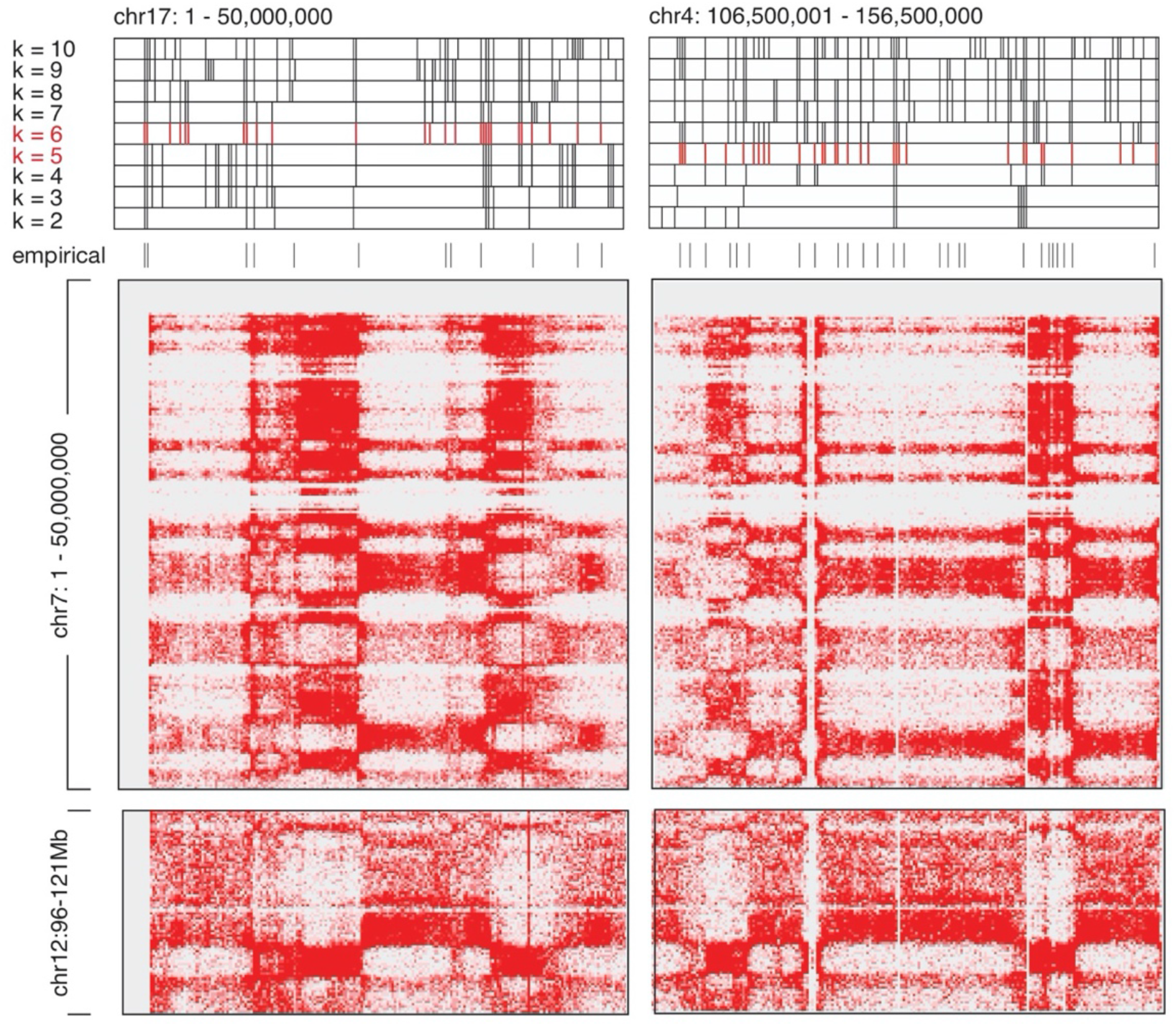
NeuN+ Hi-C clusters. Observed/expected HiC interaction matrices for NeuN+ chromatin were used to extract values using *hicdump*. K-means clustering was performed separately for odd and even chromosomes (see *Methods*). Multiple cluster numbers were tested, ranging from k=2 (bottom) to k=10 (top). Visual corroboration of cluster boundaries was performed against empirically determined boundaries (gray) based on the visual inspection of the HiC *trans* contact maps, as described in the k-means clustering strategy of Rao, et. al. (2014) (*7*). Odd chromosomes (see chr17) were most concordant with k=6 (red), while even chromosomes (see chr4) were most concordant with k=5 (red). To match and collapse these clusters across all chromosomes, any cluster comprising < 2% of the loci within the genomic space were first removed, and the remaining clusters (n=4 within odds, 4 within evens) were combined in order of size.

**Figure S2.**
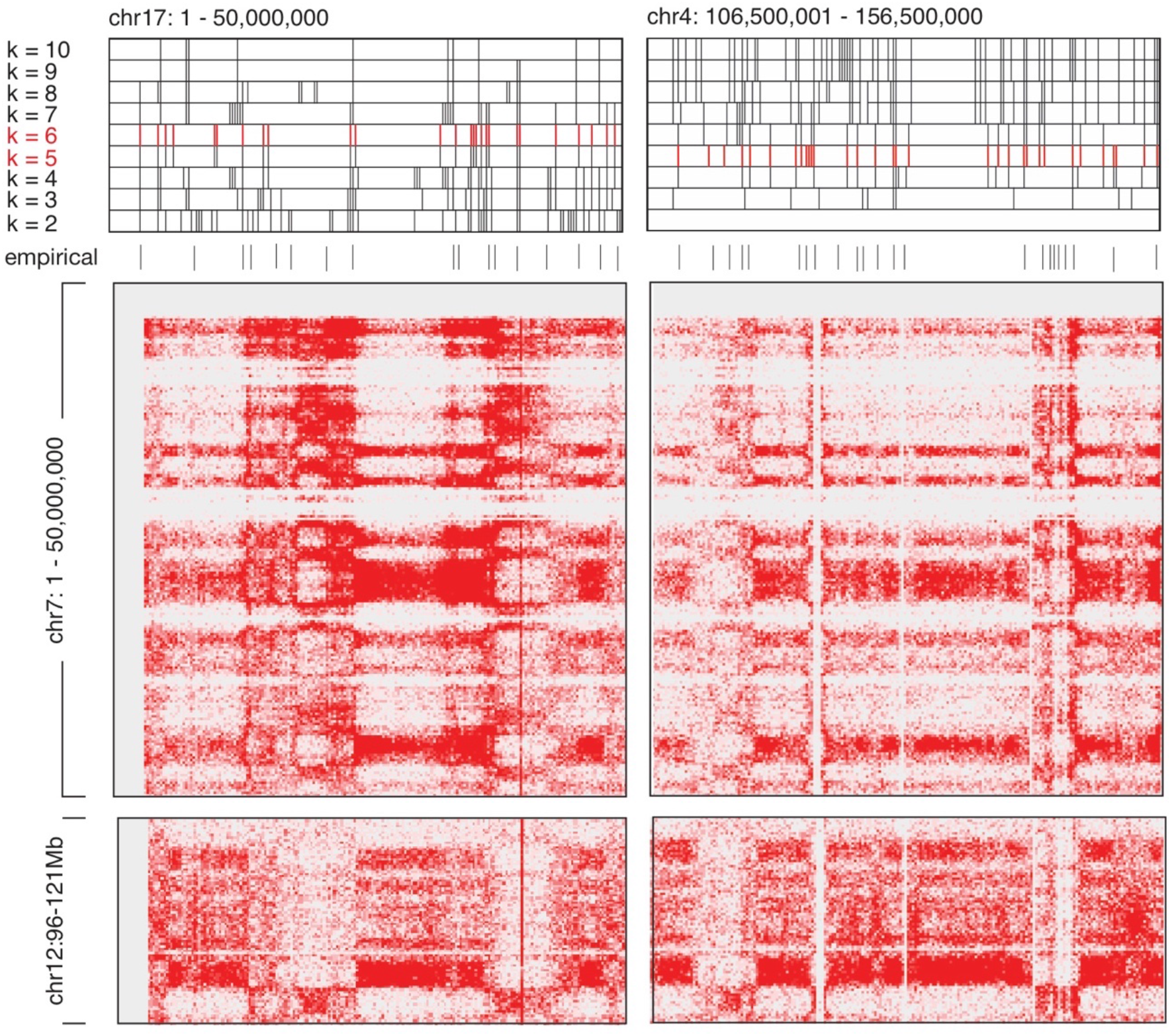
NeuN− Hi-C clusters. Observed/expected HiC interaction matrices for NeuN−chromatin were used to extract values using *hicdump*. K-means clustering was performed separately for odd and even chromosomes (see *Methods*). Multiple cluster numbers were tested, ranging from k=2 (bottom) to k=10 (top). Visual corroboration of cluster boundaries was performed against empirically determined boundaries (gray) based on the visual inspection of the HiC trans contact maps, as described in the k-means clustering strategy of Rao, et. al. (2014) (*7*). Odd chromosomes (see chr17) were most concordant with k=6 (red), while even chromosomes (see ch4) were most concordant with k=5 (red). To match and collapse these clusters across all chromosomes, any cluster comprising < 2% of the loci within the genomic space were first removed, and the remaining clusters (n=4 within odds, 4 within evens) were combined in order of size.

**Figure S3.**
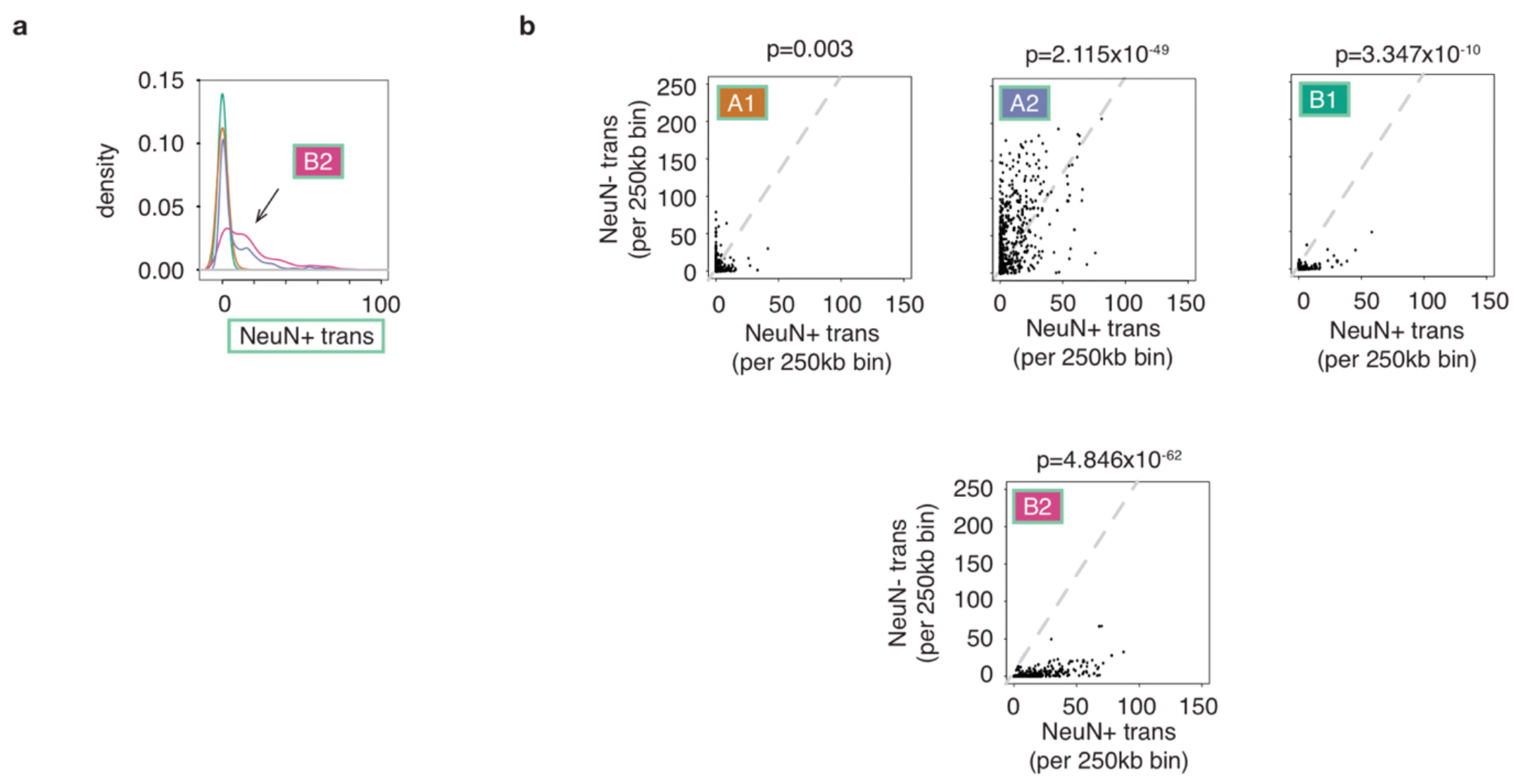
Trans interactions by subcompartment. (A) Density plot representing the NeuN+ trans-interactions occurring in 250kb bins comprising each of the four subcompartments. (B) Scatterplots of NeuN+ trans- (x-axis) vs. NeuN− trans (y-axis) for loci comprising the NeuN+ subcompartments. Dotted line in gray is the genome-wide expected line calculated from the mean trans-interactions called in NeuN+ and NeuN−.

**Figure S4.**
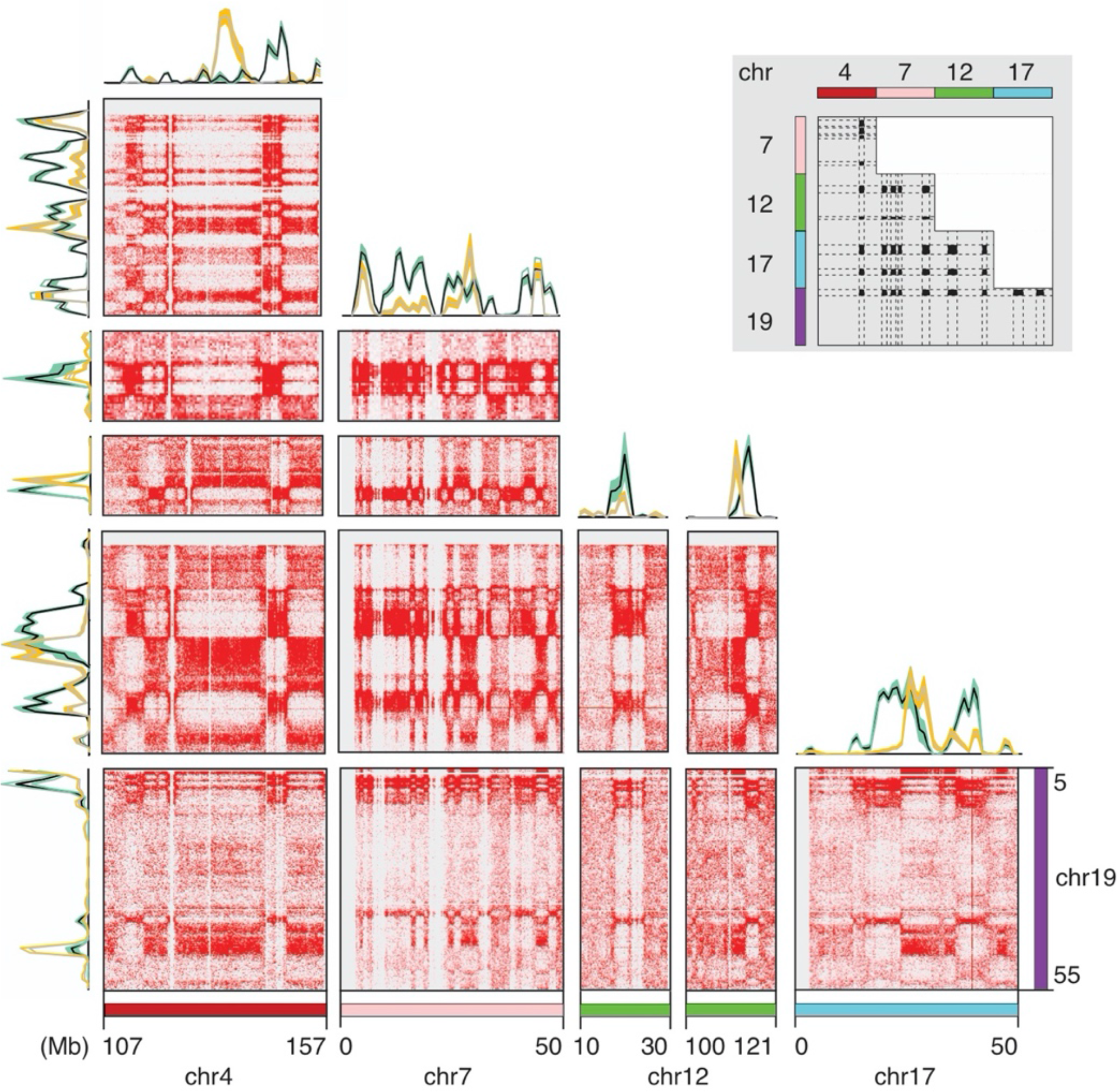
Trans interactions among B2 chromosomal megadomains. Select Hi-C pairwise matrices highlighting chromosomal megadomains of B2 subcompartment loci engaging in neuron-specific interactions in trans. Green line traces represent HOMER interactions detected in NeuN+, and orange line traces represent HOMER interactions detected in NeuN−.

**Figure S5.**
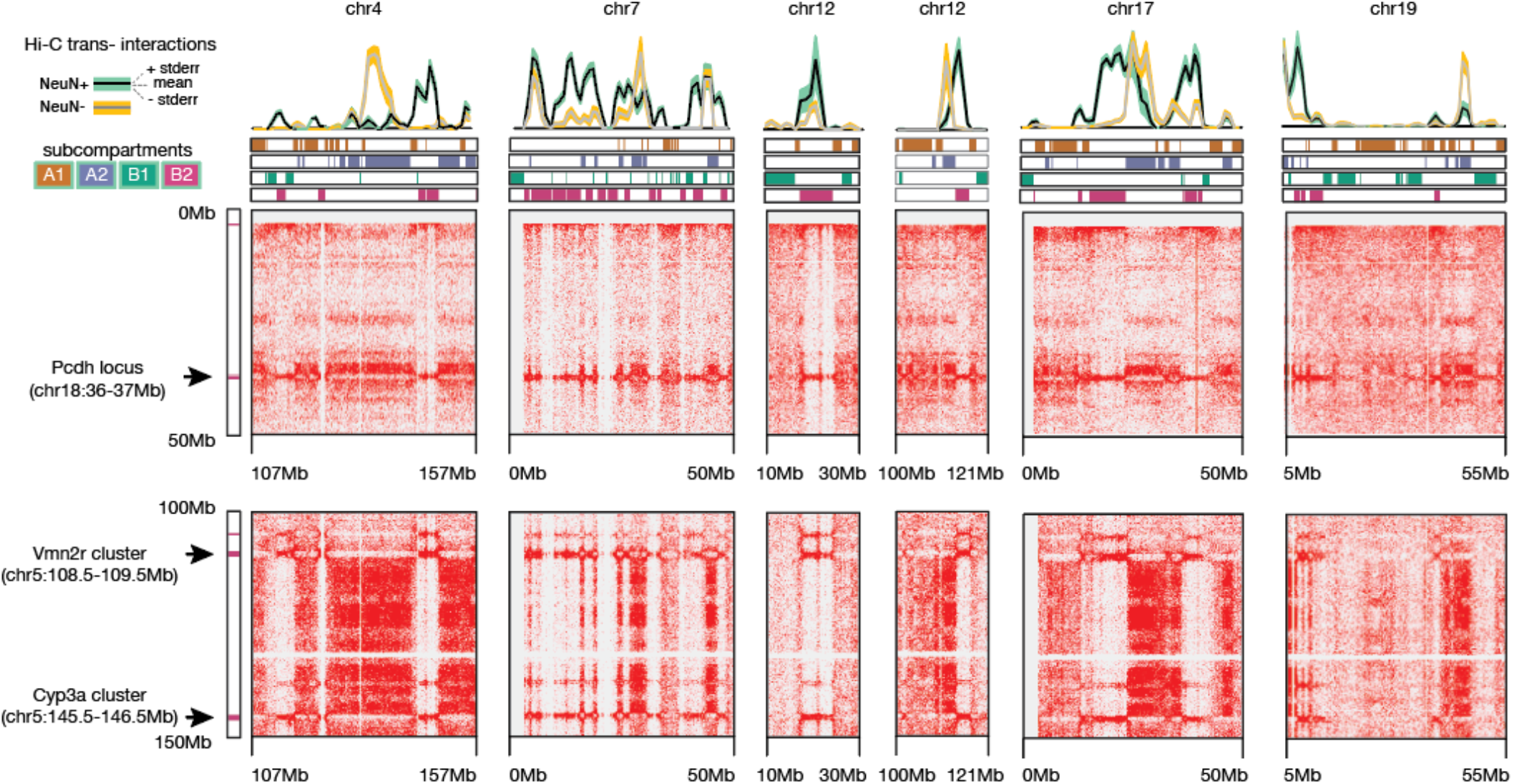
Gene clusters in B2. Hi-C pairwise contact maps encompassing multiple gene clusters (y-axis). Note coincidence of observed Hi-C interactions corresponding to each gene cluster with the B2 subcompartment loci (pink).

**Figure S6.**
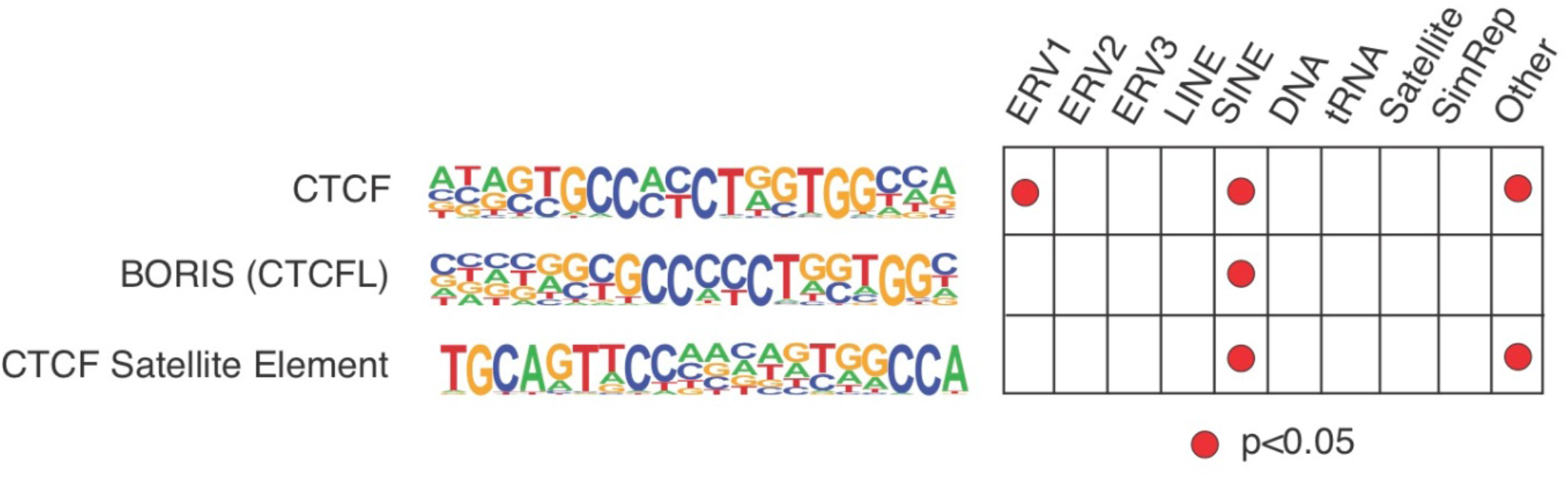
CTCF binding motif enrichment. Binding motif enrichment for CTCF and CTCF-related architectural proteins (BORIS and CTCF Satellite Element) across repetitive element categories using HOMER. Note that ERV2 do not have enrichment for CTCF and CTCF-related binding proteins, in contrast to SINE elements which are significantly associated; -log(p-values) as depicted.

**Figure S7.**
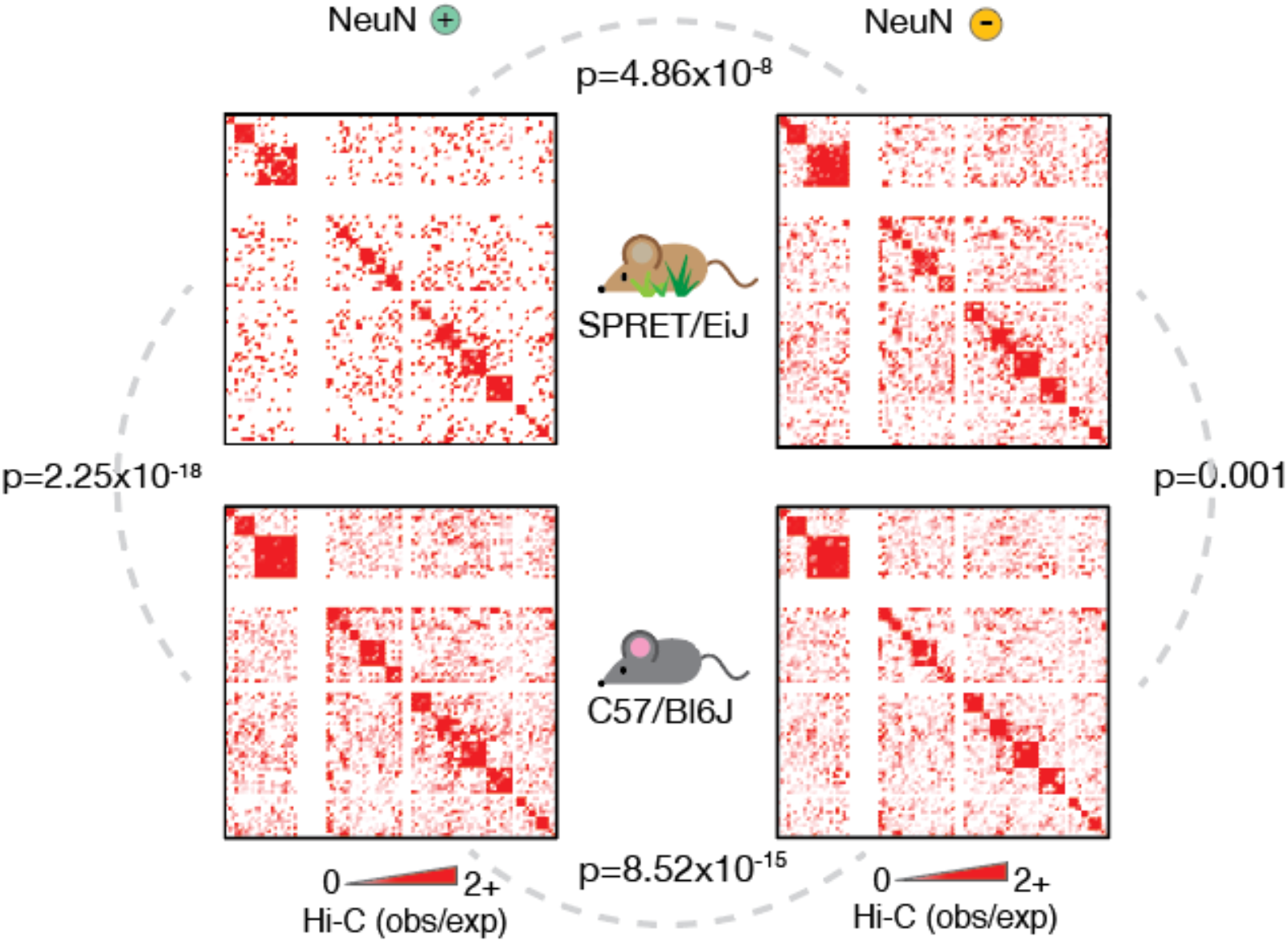
ERV2 hotspots interact least frequently in SPRET/EiJ NeuN+. Summary matrix of NeuN+ (left) and NeuN− (right) Hi-C values (observed//expected) denoting interaction frequencies between genomic loci harboring high densities of ERV2 (>99^th^ percentile, 250kb resolution) in C57/BL6J mice and SPRET/EiJ mice. Statistical significance of differences as indicated (Wilcoxon sum-rank test, two-sided, paired): SPRET/EiJ NeuN+ vs. C57/BL6J NeuN+ (p=2.25×10^−18^); SPRET/EiJ NeuN− vs. C57/BL6J NeuN− (p=0.001); SPRET/EiJ NeuN+ vs. SPRET/EiJ NeuN− (p=4.86×10^−8^); C57/BL6J NeuN+ vs. C57/BL6J NeuN− (p=8.52×10^−15^).

**Figure S8.**
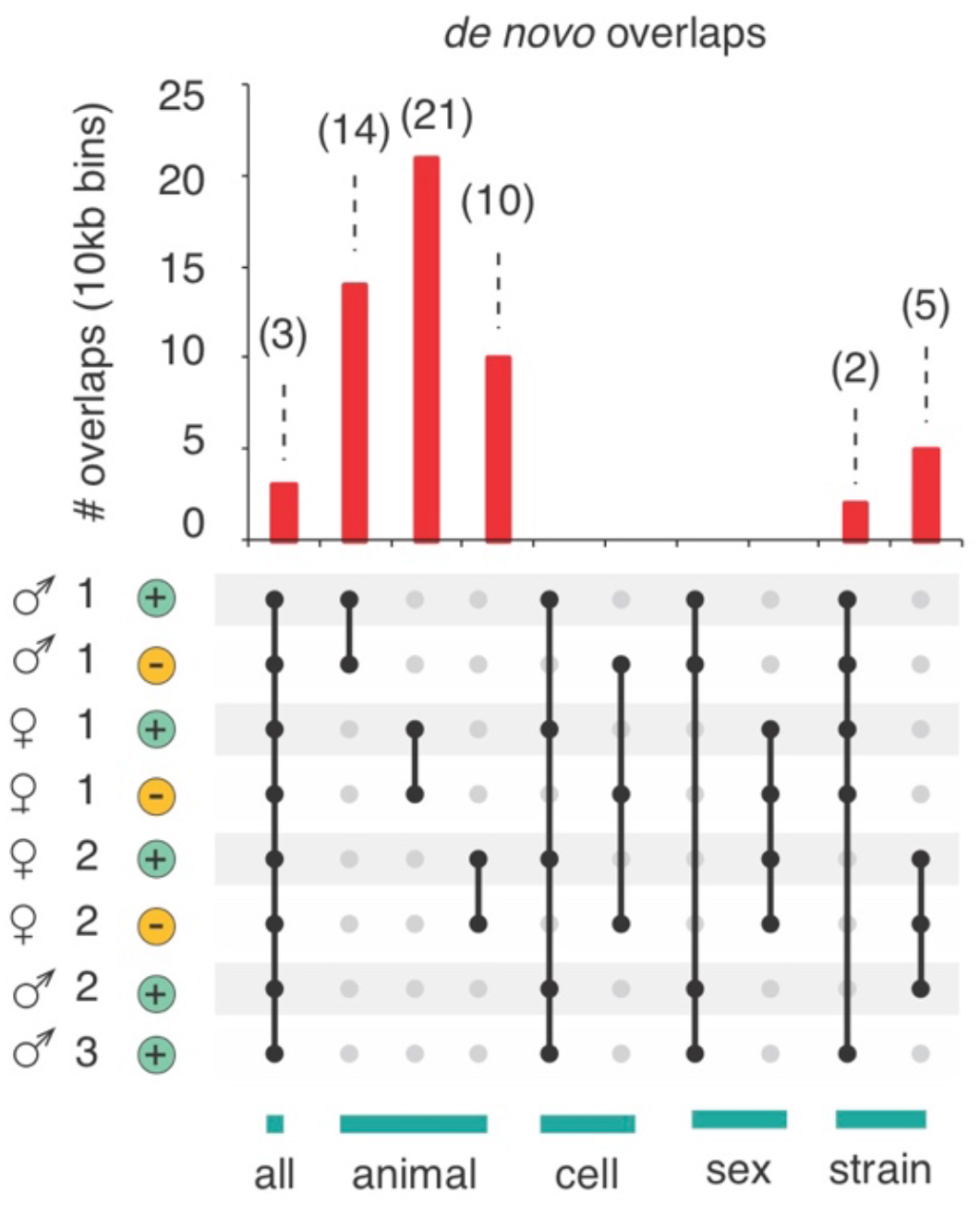
PacBio SMRT long-read sequencing of IAPEzi *de novo* integration sites. Number of overlaps (10kb resolution) shared among the different biological specimens tested (depicted as red bars). Samples are grouped by animal, cell type, sex, and strain.

**Figure S9.**
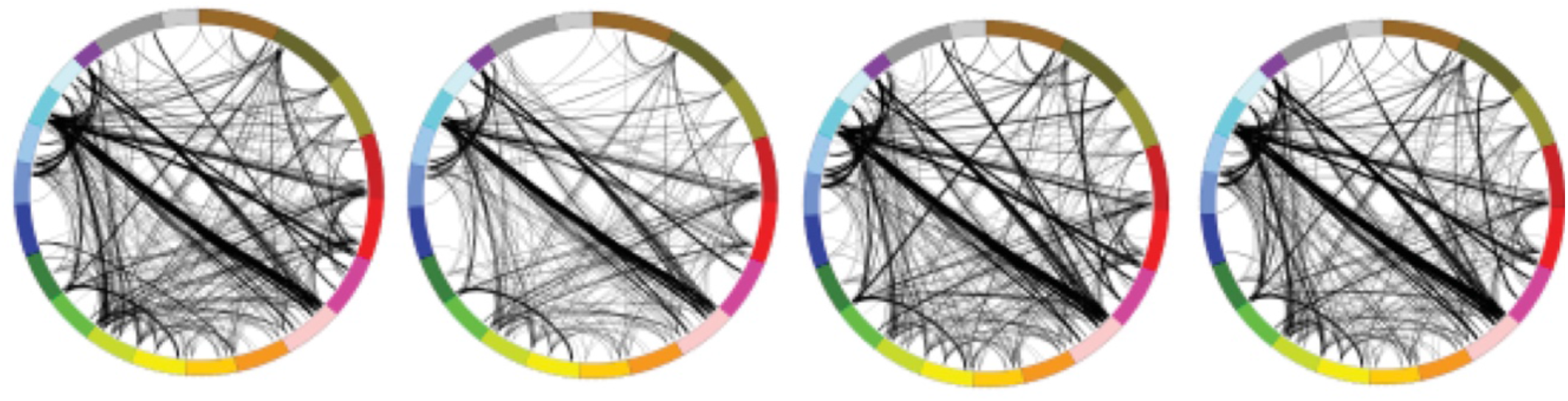
Hi-C circos plots from NeuN+ of *Camk-Cre^+^, Setdb1^2lox/2lox^* mutant mice (adult cortex). Circos plots depicting Setdb1-cKO *trans* chromosomal interactions determined by HOMER v4.8. Autosomes (chr1- 19) and chrX/Y are depicted along the periphery of the circle. Significance threshold: p < 10^−50^.

**Figure S10.**
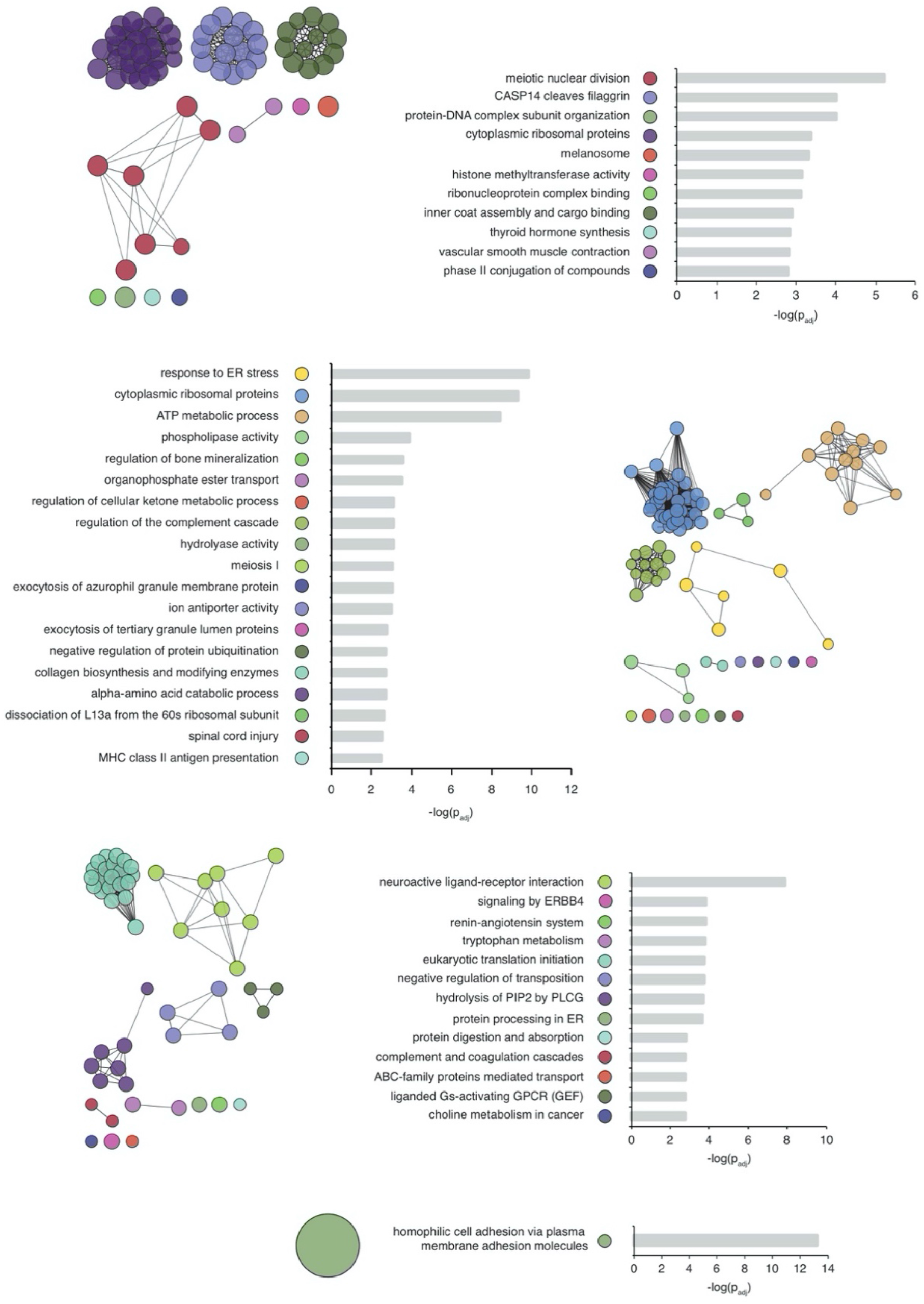
GOs for elevated transcripts in *Camk-Cre^+^, Setdb1^2lox/2lox^* mutant vs. control adult cortex, divided by subcompartment. Top ten categories displayed, with corresponding enrichment significance for each GO category (-log(p-adj)) (ClueGO (v2.5.1) for Cytoscape (v3.6.1, right-sided hypergeometric test analysis with Benjamini-Hochberg p-value correction)). Gene modules depicted with connected nodes.

**Figure S11.**
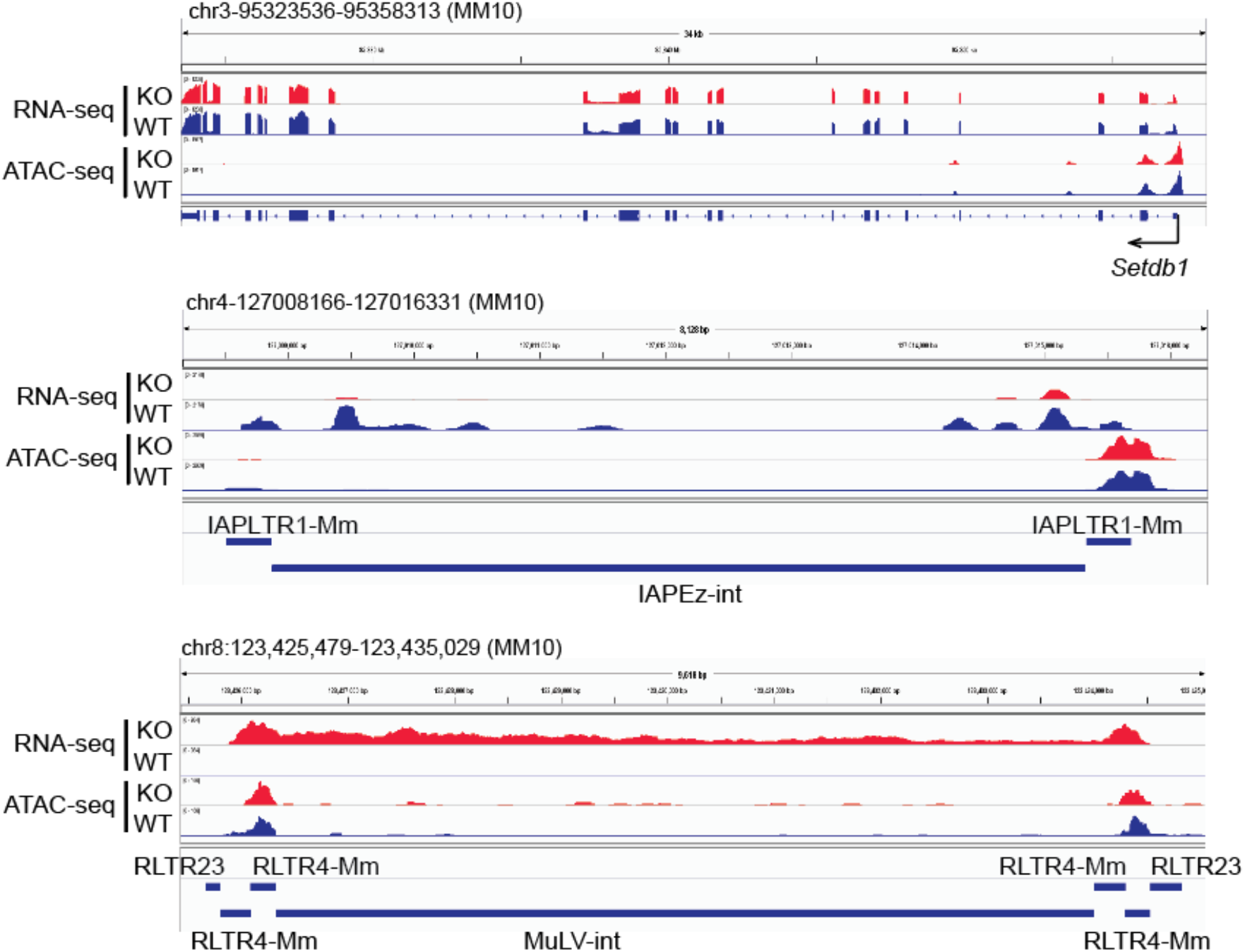
Microglia RNA-Seq and ATAC-Seq. Representative images show signal at (Top) *Setdb1* gene, (Middle) IAPEz-int, and (Bottom) ERV1-MuLV loci from microglia-specific RNA-seq and ATAC-seq in *Setdb1- CK-cKO* and controls. Notice no change for *Setdb1* transciption, especially exon III that was ablated in our neuronal knockout system. And also, no change for the chromatin accessibility at the locus. Notice the decrease of IAPEz-int and increase of MuLV transcripts at these two specific loci.

**Figure S12.**
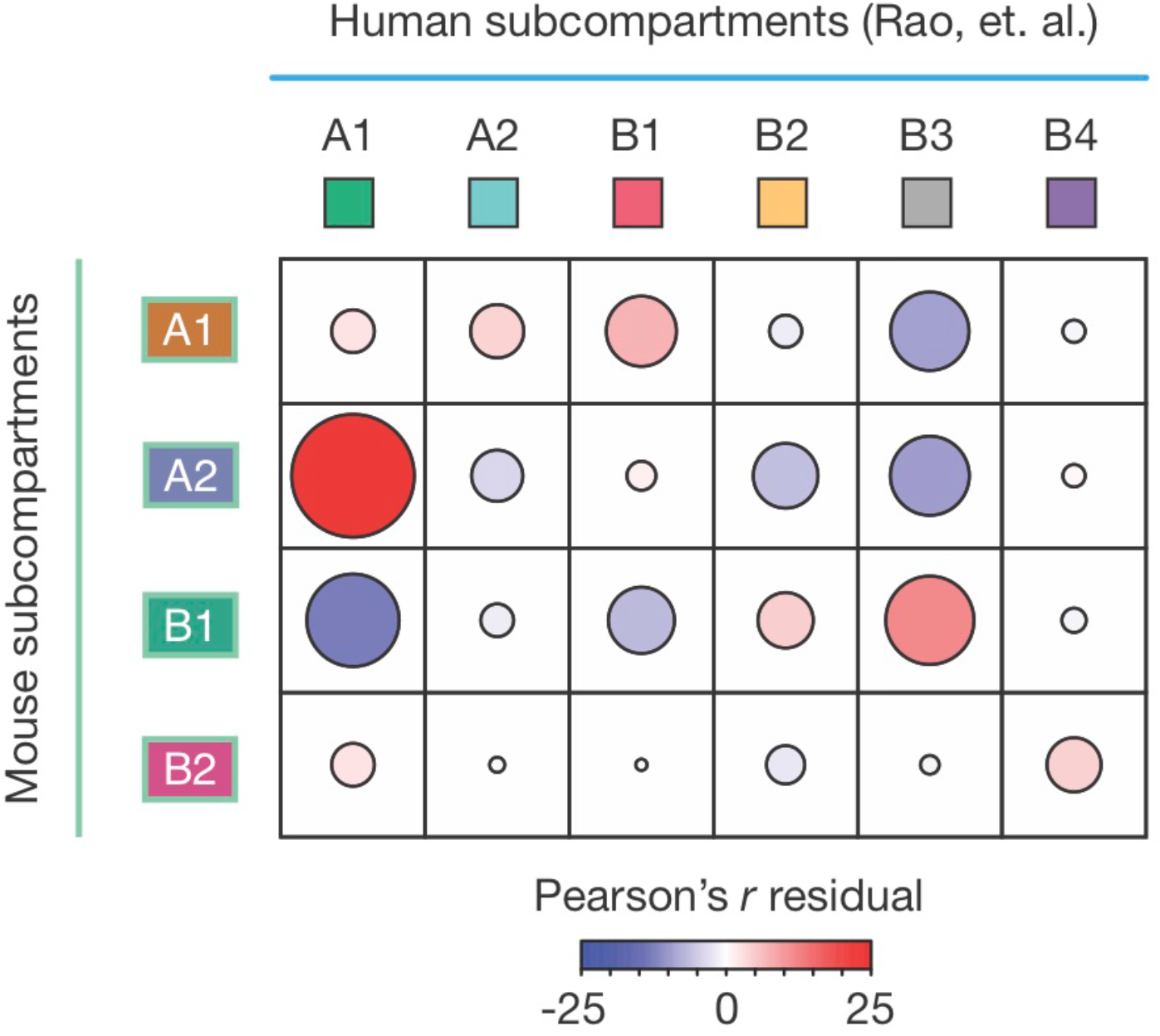
Mouse/human subcompartment comparisons. Observed/expected (Pearson’s residual r) length of overlaps for lifted over coordinates from the human subcompartments (Rao, et. al. (2014)) (*7*) with the NeuN+ subcompartments. The size of each ellipse represents the |r|_abs_, and the color of each ellipse represents r.

**Figure S13.**
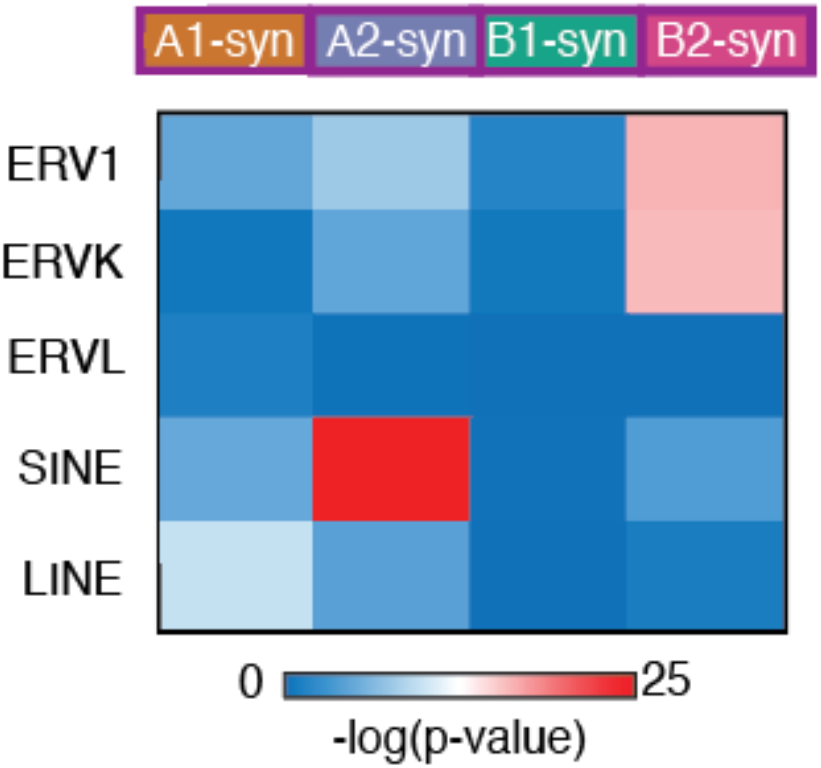
Inter-chromosomal contacts in human neurons are enriched with (H)ERV retroelements. Heatmap of significance of genome-wide associations between syntenic genomic loci for each of the mouse subcompartments and repetitive element hotspots in humans.

